# The Neural Basis for a Persistent Internal State in *Drosophila* Females

**DOI:** 10.1101/2020.02.13.947952

**Authors:** David Deutsch, Diego A. Pacheco, Lucas J. Encarnacion-Rivera, Talmo Pereira, Ramie Fathy, Adam Calhoun, Elise C. Ireland, Austin T. Burke, Sven Dorkenwald, Claire McKellar, Thomas Macrina, Ran Lu, Kisuk Lee, Nico Kemnitz, Dodam Ih, Manuel Castro, Akhilesh Halageri, Chris Jordan, William Silversmith, Jingpeng Wu, H. Sebastian Seung, Mala Murthy

## Abstract

Sustained changes in mood or action require persistent changes in neural activity, but it has been difficult to identify and characterize the neural circuit mechanisms that underlie persistent activity and contribute to long-lasting changes in behavior. Here, we focus on changes in the behavioral state of *Drosophila* females that persist for minutes following optogenetic activation of a single class of central brain neurons termed pC1. We find that female pC1 neurons drive a variety of persistent behaviors in the presence of males, including increased receptivity, shoving, and chasing. By reconstructing cells in a volume electron microscopic image of the female brain, we classify 7 different pC1 cell types and, using cell type specific driver lines, determine that one of these, pC1-Alpha, is responsible for driving persistent female shoving and chasing. Using calcium imaging, we locate sites of minutes-long persistent neural activity in the brain, which include pC1 neurons themselves. Finally, we exhaustively reconstruct all synaptic partners of a single pC1-Alpha neuron, and find recurrent connectivity that could support the persistent neural activity. Our work thus links minutes-long persistent changes in behavior with persistent neural activity and recurrent circuit architecture in the female brain.

## Introduction

Internal brain states influence perceptions, decision-making, and actions (Anderson, 2016; Berridge, 2004; Lorenz and Leyhausen, 1973) - for example, when hungry, we make different decisions about what to do (prioritizing searching out food over other tasks) and what to eat (expanding the repertoire of foods we find appetizing) based on our hunger status (Sayin et al., 2019; Sternson et al., 2013). The timescales of persistence of an internal brain state can vary - some, like sleep-wake cycles, persist over hours (Scammell et al., 2017; Shaw et al., 2000), while others may last only a few seconds (Calhoun et al., 2019). During social interactions, internal state may correspond to levels of arousal or drive (Anderson, 2016; Berridge, 2004; Lorenz and Leyhausen, 1973), and can impact whether and how individuals interact, with consequences for mating decisions and reproduction (Chen and Hong, 2018; Kennedy et al., 2014; Stowers and Liberles, 2016). In general, social internal states have been studied by activating small subsets of neurons, and examining behaviors, such as courtship or aggression, that outlast the activation of these neurons.

In flies, a small population of male-specific neurons (P1) that express the sex-specific transcription factors Fruitless and Doublesex (Auer and Benton, 2016), drive both male-aggression and male-mating behaviors (Hoopfer et al., 2015; Koganezawa et al., 2016; von Philipsborn et al., 2011). P1 neurons are a subset of the larger Doublesex+ pC1 neural subset (Kimura et al., 2008). Brief optogenetic activation of P1 neurons drives both persistent song production in solitary males and persistent aggression upon introduction of another male, both over minutes (Bath et al., 2014; Hoopfer et al., 2015; Inagaki et al., 2014). The specific timescales of persistency for these two behaviors scales with different overall levels of P1 activity. While P1 activation is sufficient for eliciting the persistent behavioral phenotypes, other groups of neurons are involved in maintaining the persistent state (Jung et al., 2020; Zhang et al., 2019). Whether P1 neurons themselves are (Zhang et al., 2018) or are not (Inagaki et al., 2014) active during the persistent period may depend on the details of the stimulation protocol. Work on P1 in flies bears some similarity to work in mice. In male mice, optogenetic activation of SF1 (steroidogenic factor 1) expressing neurons in the dorsomedial part of the ventromedial hypothalamus (VMHdm^SF1^) drive multiple defensive behaviors (Wang et al., 2015), that outlast the stimulation by up to one minute (Kunwar et al., 2015; Wang et al., 2015). Fiber photometry in VMHdm^SF1^ neurons expressing GCaMP in freely behaving mice exposed to an anesthetized rat, revealed activity that persisted for over a minute following rat removal (Kennedy et al. 2019), but the circuit mechanisms that support this persistent activity are not yet known.

Little attention has been paid to the persistence of social behaviors in females. *Drosophila* females lack P1 neurons, but do have Doublesex+ pC1 neurons (Rideout et al., 2010; Robinett et al., 2010; Zhou et al., 2014), including a subset that are female-specific (Wu et al., 2019). Silencing and activation of pC1 neurons can affect female receptivity (Zhou et al., 2014), making these neurons candidates (similar to P1 neurons in males) for driving a persistent internal state. Thermogenetic activation of pC1 neurons in females drives chasing (Rezával et al., 2016; Wu et al., 2019), a male-specific behavior, and also aggressive behaviors toward females (Palavicino-Maggio et al., 2019). Taken together these data suggest that female pC1 neurons can drive multiple distinct behaviors, similar to male P1 neurons, but whether female pC1 neurons can drive persistent changes in behavior and persistent neural activity, has not yet been investigated.

Understanding the control of internal states in females is important for at least two reasons. First, as males and females produce distinct behaviors during courtship and mating, it is critical to address which aspects of the circuit are shared between sexes, and which are sex-specific (Deutsch et al., 2019; Kohl et al., 2013). Second, as the female makes the ultimate decision – to mate or not to mate - examining how her internal state shapes her responses to the male’s cues and her mating decisions has clear ethological relevance. Here, we investigate the circuit mechanisms underlying a persistent internal state in females. We find that pC1 activation drives persistent changes in female behavior for minutes following stimulus offset. The effect of pC1 activation on female receptivity peaks at a different time relative to stimulus offset, compared with effects on aggressive and male-like behaviors. We identify the subset of pC1 neurons (called pC1-Alpha (Wu et al., 2019)) that affects the persistent aggressive and male-like behaviors. Using calcium imaging, we find that pC1-Alpha activation can elicit persistent activity among multiple cell types, including pC1 neurons themselves. Finally, we leverage the automated segmentation of a complete EM volume of the female brain (Zheng et al., 2018), to map all inputs and outputs of a pC1-Alpha neuron and uncover a putative circuit basis for the persistent internal state.

## Results

### Female pC1 activation persistently modulates both female receptivity and song responses

To investigate the neural basis of a persistent internal state in the female brain, we focused on pC1 neurons, one of eight Doublesex-expressing cell types in the central brain (Fig. 1A; (Kimura et al., 2015)). We used an intersection between two driver lines (Dsx-GAL4 and R71G01-LexA (hereafter referred to as pC1-Int; see Supplemental Table 1 for list of genotypes used in this study); Fig. S3E), to label pC1 neurons, as done previously (Rezával et al., 2016; Zhou et al., 2014). We tracked male and female body parts (head and thorax) in addition to recording all sounds (Fig. 1B, Supp Fig. S1A, Supp Movie S1; see Methods for details on song segmentation and tracking of flies on a non-homogenous background). Silencing pC1-Int neurons in females had previously been shown to affect receptivity (Zhou et al., 2014); we corroborated these results (Fig. 1C) and additionally showed that constitutive silencing of pC1-Int neurons diminished responses to male song (Fig. 1D). To identify persistent changes in behavior following pC1 activation, we developed a new paradigm in which we activated pC1-Int in a solitary virgin female for 5 minutes, followed by a variable delay period, after which a virgin male was introduced to examine female behaviors in the context of courtship (Fig. 1E) - there was no optogenetic activation following the first 5 minutes. The activity of stimulated neurons should decay during the variable delay period (d0 (0 minute delay), d3 (3 minute delay), or d6 (6 minute delay)) - we test this explicitly below. Therefore, the effects of differing levels of persistent activity on behavior could be uncovered.

**Figure 1:**
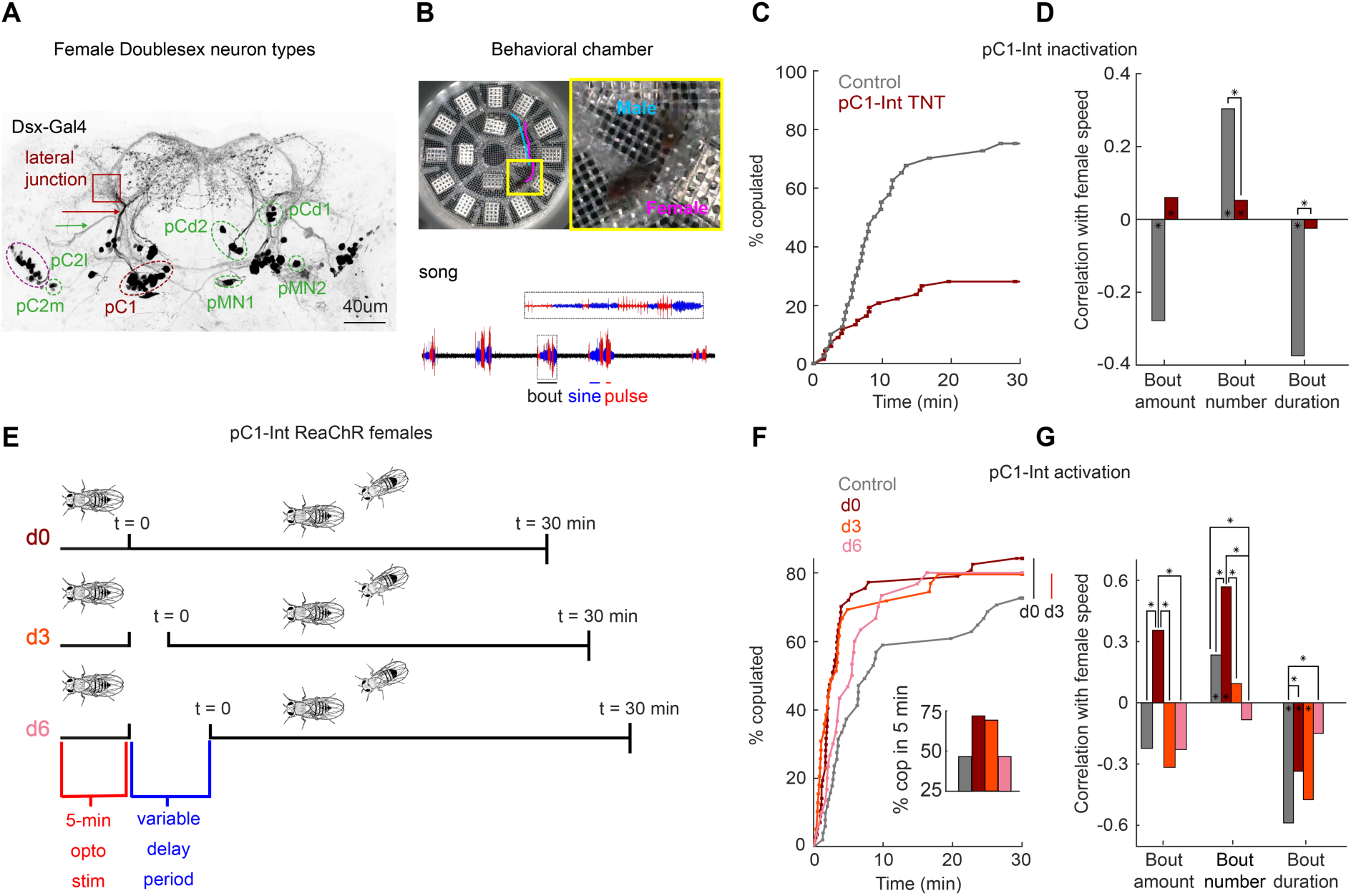
pC1-Int activation has a persistent effect on female receptivity and responses to male courtship song. **(A)** Anatomy of Dsx+ neurons in the female brain. Max z-projection of a confocal stack of a fly brain in which Dsx+ cells are labeled with GFP (adapted from (Deutsch et al., 2019)). Dsx is expressed in 8 morphologically distinct cell types in the female brain (7 types are indicated by circling of their somas; the more anterior cell type aDN (Lee et al., 2002) is not shown). pCd has two morphologically distinct types, pCd1 and pCd2 (Kimura et al., 2015). Many of these cells project to a brain region known as the lateral junction (red square). pC1 cells project to the lateral junction through a thin bundle (marked with red arrow), and pC2l (but not pC2m) cells project to the contralateral hemisphere through a unique bundle (green arrow). These bundles are used for identifying the cells at the EM dataset (see Methods). **(B)** The behavioral chamber (diameter ∼25mm) is tiled with 16 microphones and fitted with a camera above to record fly movements. Male and female positions were tracked offline (see Methods; a 1.5 second example trace is shown for the male (cyan) and female (magenta)). Male song was automatically segmented into sine and pulse song. An example song trace (1.5 seconds) is shown, zooming on a single song bout (black dotted box). **(C)** Percent of male/female pairs that copulated as a function of time. Copulation rate was lower when a female expressed TNT in pC1-Int cells (Cox proportional hazards regression, see Methods; dark red, N = 68 pairs; P = 6.3*10^-6^) compared with controls (grey, N = 40 pairs). pC1-Int TNT: R71G01-LexA/ UAS>STOP>TNT; LexAop2-FLP; Dsx-Gal4. **(D)** Rank correlation between male song (bout amount, number and duration) and female speed (see Methods for definitions of song parameters and correlation calculation) for TNT expressing females (dark red) or controls (grey). Significance (indicated by an asterisk above the line connecting a pair of groups), was measured using ANOCOVA (MATLAB aoctool) and multiple comparison correction (*P<0.01). An asterisk in the base on a bar indicates a significant correlation between a single male song measure and female speed (MATLAB corr, *P<0.01). **(E)** Experimental design for pC1-Int activation. pC1-Int cells were activated (using ReaChR) for 5 minutes in a solitary female placed in the behavioral chamber. Following light offset, a wild type male was introduced at t=0, with a variable delay period (d0 = no delay; d3 = 3min delay; d6 = 6min delay) prior to introduction of the male. All behavioral phenotypes were measured following pC1-Int activation at t > 0. pC1-Int ReaChR: R71G01-LexA/LexAop2-FLP;Dsx-Gal4/UAS>STOP>ReaChR. **(F)** Same as (C), but for females expressing ReachR in pC1-Int cells according to the protocol shown in (E). Inset: The percent of pairs copulated between t = 0 and t = 5 minutes for each condition. pC1-Int activated females in the d0 condition (N = 57) copulated significantly faster than controls (N = 51; vertical black line; P = 0.0045, Cox’s proportional hazards regression model, accounting for censoring, as not all flies copulated in 30 minutes; black vertical line). Time to copulation was also shorter in the d3 group (N = 39 pairs) compared with controls (d0, no ATR), but the difference was not significant after Bonferroni correction (P = 0.034; red vertical line), and no significant difference was found between the 6-min delay (N = 30) and control groups (P = 0.21). **(G)** Same as (D), but for females expressing ReachR in pC1-Int cells according to the protocol shown in (E). Asterisks show significance, using the same criteria as in (D). Numbers of pairs are the same as in (F).

Experimental flies were fed all-trans-retinal (ATR), which is required for ReaChR (red-shifted Channelrhodopsin) function in flies (Inagaki et al., 2014)). Control flies shared the same genotype but were not fed ATR. We found that activation of pC1-Int neurons induces a persistent effect on female receptivity and responses to male song, but that this effect diminishes with a delay period between neural activation and introduction of a male. pC1-Int activated females copulated significantly faster than controls in the d0 condition, with reduced copulations following a delay between optogenetic activation and introduction of a male (Fig. 1F). About 75% of the flies copulated within 5 minutes in the d0 and d3 conditions, compared with fewer than 50% in the control and d6 groups (Fig. 1F, inset). Activating pC1-Int also produced a persistent effect on responses to male song, overall with the strongest effect at the d0 delay (Fig. 1G) - d0 females, in comparison with controls, accelerated more in response to all song elements, behaving like unreceptive females (Coen et al., 2014). This effect was not due to changes in wild type male song structure for the d0 condition, although male song bouts were shorter and less frequent in the d3 and d6 conditions (Supp Fig. S1B), possibly due to the strong effect of pC1-Int activation on male-female interactions at longer delays, as shown below. These results suggest that the effect of pC1-Int activation on female receptivity and female song responses likely occurs via parallel downstream pathways. Another set of Dsx+ neurons called pCd1 (Fig. 1A) was shown to control female receptivity (Zhou et al., 2014) and to enable P1 induced persistent activity in males (Jung et al., 2020). While pCd1 silencing in females reduced receptivity (Supp Fig. S1C), pCd1 activation had no persistent effect on female receptivity (Supp Fig. S1D, left). pCd1 activation affected female responses to male song (Supp Fig. S1D, right), further supporting the conclusion that these behaviors (receptivity and song responses) are controlled via separate pathways.

### Female pC1 activation drives persistent female shoving and chasing

To quantify other behaviors elicited by pC1-Int activation, we decomposed male and female movements and interactions into 17 parameters (Calhoun et al., 2019) (Fig. 2A). We used a Support-Vector Machine (SVM) framework (Cortes and Vapnik, 1995; Cristianini et al., 2000) to find the weights that best classify single frames as belonging to sessions of control vs experimental groups (all delay conditions, d0-d6). We found that the weights of 8 out of 17 parameters were significantly different from zero (Fig. 2B and Supp Fig. S1E), with the strongest weight being fmAngle, defined as the degrees the female needs to turn in order to point toward the male centroid. The weight of fmAngle is negative because this parameter is smaller in the experimental flies compared with controls, indicating that pC1-Int activated females spend more time facing the male (Supp Fig. S1F). When separated by experimental condition, the SVM classifier performed best on the d3 condition versus control (Fig. 2C), indicating that male-female movements and interactions are most distinct following this delay.

**Figure 2:**
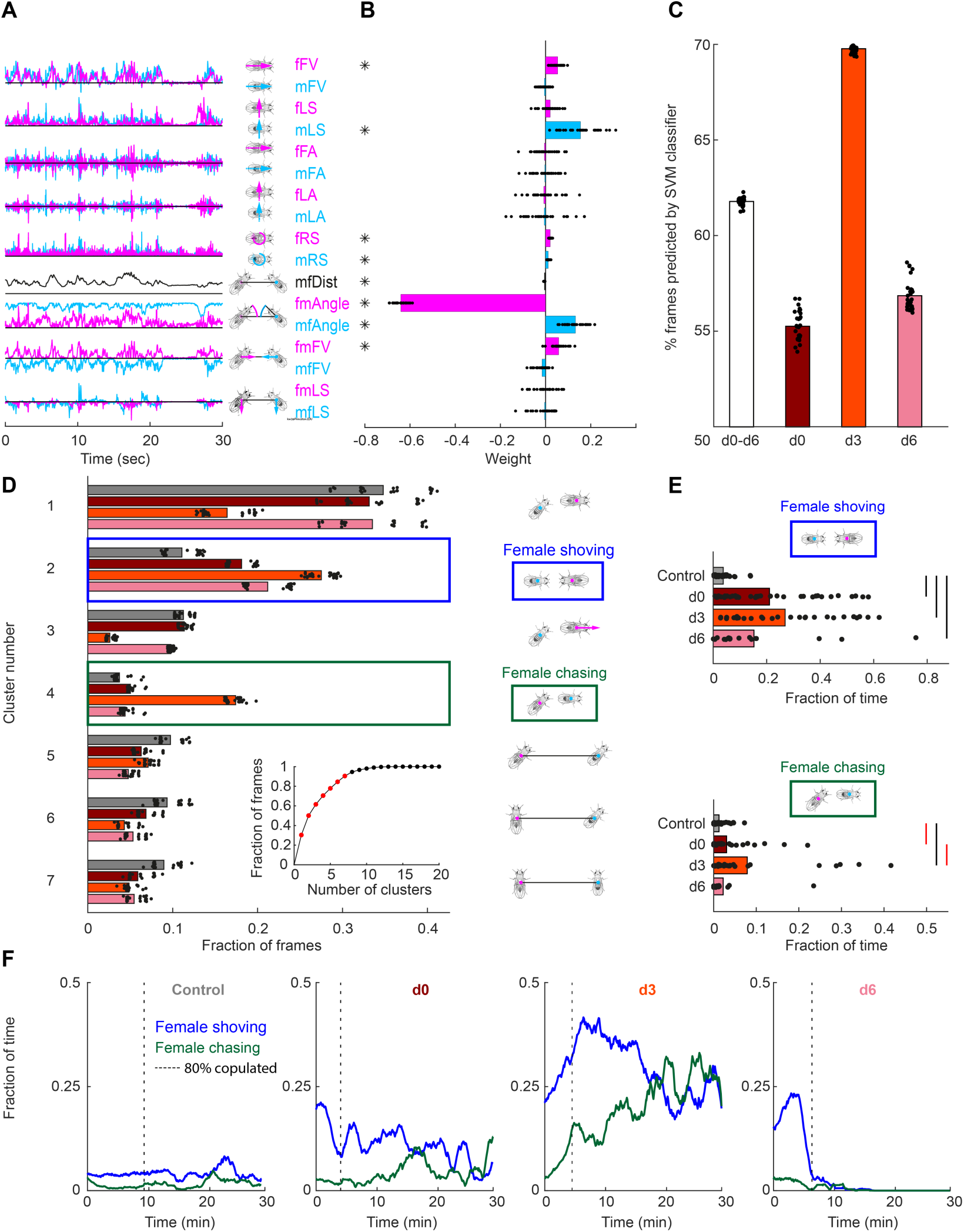
Automated identification of persistent female behaviors following pC1-Int activation. **(A)** For each video frame, 17 parameters were extracted based on the tracking of male/female position and heading (see Methods for parameter definition). An example trace (30 seconds) is shown for each parameter. **(B)** 30 independent (using non-overlapping sets of video frames) Support Vector Machine (SVM) classifiers were trained to classify frames (each frame represented by 17 values) as belonging to control or experimental group (all delays d0-d6 considered together; see Methods). Each classifier is represented by 17 points-one for each parameter. Each point is the weight associated with a given parameter for one classifier, and the bar height represents the mean over classifiers (* P<10^-4^, one-sample t-test; see Methods). **(C)** The percent of frames correctly classified using the SVM classifier. Each dot is the prediction of a single SVM classifier, trained to classify frames as belonging to control or experimental group (do, d3, d6 or d0-d6 taken together). The bar is the mean prediction over 30 classifiers, for one group. **(D)** K-means was used to cluster frames based on the 8 most significant parameters (marked with asterisks in (B)). The largest 7 clusters include 90.4% of the frames (see inset). Clustering was performed 30 times (black dots), using different but overlapping sets of frames. The same number of frames was taken from each group (see Methods). Bar height indicates the average over all 30 repeats from a given group and cluster. Cluster 2 (blue box - ‘female shoving’) is more probable following pC1-Int activation (in both d0, d3 and d6 conditions) compared to control, while cluster 4 (green box-‘female chasing’) is more probable in the d3 condition only compared to control. Cartoon on the right, for each cluster, is a schematic describing the male-female interaction, based on the mean values of the weights. **(E)** JAABA based classification of shoving (top) and chasing (bottom) behaviors. Each dot represents a single pair of flies. The fraction of time the male-female pair spent shoving (0.037/0.21/0.27/0.15 for control/d0/d3/d6) or chasing (0.013/0.030/0.079/0.022) are shown. Black lines represent significant differences with p<0.05 after Bonferroni correction for multiple comparison. Red lines - significant before, but not after correction for multiple comparisons. **(F)** Fraction of time the female spent chasing or shoving (moving average with a two-minute window; mean over all flies in a given condition), based on JAABA classification in each condition (control, d0, d3, d6). T = 0 is the time the male was introduced (see Fig. 1E), and the vertical dotted line indicates the time, for each condition, when 80% of the pairs copulated.

Next, we clustered individual video frames based on the values of the 8 parameters identified as most important by the SVM (Supp Fig. 1E, asterisks), and found that the largest 7 clusters accounted for over 90% of all frames (Fig. 2D, inset). The weights of the 8 parameters were different for each cluster (Supp Fig. S1G), representing different behaviors. Five of the clusters describe behaviors that are the same or reduced following pC1-Int activation (clusters 1 and 3, male chasing female, and 5-7, increased male-female distance). Two clusters, however, describe behaviors that occur with higher probability following pC1-Int activation (clusters 2 and 4; Fig. 2D). Cluster 2 is characterized by small fmAngle and small mfAngle (indicating that the male and female are facing each other), decreased male-female distance, and large fmFV (indicating that the female is close to the male and moving in his direction). Cluster 4 is characterized by small fmAngle and large mfAngle (indicating that the female is behind the male), decreased male-female distance, and large fmFV (indicating that the female is moving in the direction of the male). Based on the weight values, and verified by inspection of the videos following clustering (Supp Movie S2), we termed cluster 2 ‘female shoving’ and cluster 4 ‘female chasing’. For both female shoving and female chasing, the amount of each behavior was highest in the d3 condition relative to control (Fig. 2D).

We then used JAABA (Kabra et al., 2013) to train a classifier to recognize epochs (groups of video frames) of female chasing and female shoving in the data (Supp Fig. 1H; see Methods). This analysis confirmed that the amount of female shoving and chasing was greatest in the d3 condition relative to control (Fig. 2E), in addition to revealing that the duration of female chasing and shoving bouts was longer (Supp Fig. 1I). Female shoving and chasing persisted for as long as 30 minutes in the d3 condition (Fig. 2F), but not in the d0 and d6 conditions, indicating that cues from the courting male contributed to the persistent behaviors. While the percent of time the female spent shoving or chasing in the first two minutes after the male was introduced was similar in the d0 and d3 conditions (19.7/21.3% for shoving, 2.8/3.1% for chasing), shoving and chasing probabilities rose over time in the d3, but not in the d0 condition. In the d6 condition, shoving probability was comparable to the probability in the d0-d3 conditions in the first two minutes (15%), but decayed to control level after 6 minutes. These data suggest a difference in female brain state between these three conditions, differences that impact her interactions with a male.

Manual inspection of the videos confirmed the results above (Fig. S2B), and in addition, identified several additional behaviors produced by females following pC1-Int activation. These include ‘female approaching’, ‘circling’, ‘head-butting’, and ‘female wing extension’ (Supp Fig. S2C-F; Supp Movies 2,3).

We found that some of these behaviors were coupled; for example, ‘circling’ was often preceded by ‘female shoving’ (Supp Fig. S2D and Supp Movie S3) and ‘female wing extension’ was often coincident with ‘female chasing’ (Supp Fig. S2G and Supp Movie S2), similar to male behavior during courtship (although we did not observe sounds from the females that resembled male courtship song (Supp Fig. S2F)). Our automated classifier (Fig. 1B) did not find these behaviors because we only tracked the head and thorax of each fly, which did not provide enough information to automatically identify these behaviors, or to keep accurate track of identities during behaviors in which the male and female often overlap (e.g., during ‘circling’). We observed no female shoving or chasing following pCd1 activation (Supp Fig. S2A).

In sum, we found that for minutes following pC1-Int activation, females produced a variety of behaviors directed at the male. Some of these appear aggressive, such as shoving and head-butting (Nilsen et al., 2004; Palavicino-Maggio et al., 2019), while others resemble male courtship behaviors, such as chasing and unilateral wing extension. These behaviors typically peaked in the d3 condition, where they remained high 30 minutes after male introduction. In contrast, the effect on female receptivity and female responses to male song, were both strongest in the d0 condition. Below we further explore the timescales of these behaviors.

### pC1 cell types

We propose three possible circuit configurations to explain our behavioral results (Fig. 3A). In the first configuration, the same set of pC1 neurons activate different downstream circuits, each one controlling a different behavior. The differences in the temporal dynamics arise downstream of pC1. In the second configuration, three non-overlapping subsets of pC1 neurons control the different behaviors. In the third configuration, one pC1 subset controls female receptivity (that peaks at d0), and another set controls chasing and shoving (both peaking at d3). The second and third models assume some heterogeneity in the pC1 population. To evaluate these circuit models we examined the behavioral consequences of activating distinct subsets of pC1 neurons. To define pC1 cell types, we used automated reconstruction of neurons in an EM volume of a female brain (FAFB) (Zheng et al., 2018); neuron segmentation and reconstruction was accomplished using a novel platform for visualization and proofreading (flywire.ai, manuscript in preparation). We examined the morphologies of neurons that send projections to the lateral junction through a thin neuronal bundle similar to known pC1 neurons (Fig. 1A, red arrow; Fig. 3B, red line; see also (Deutsch et al., 2019; Zhou et al., 2014).

**Figure 3:**
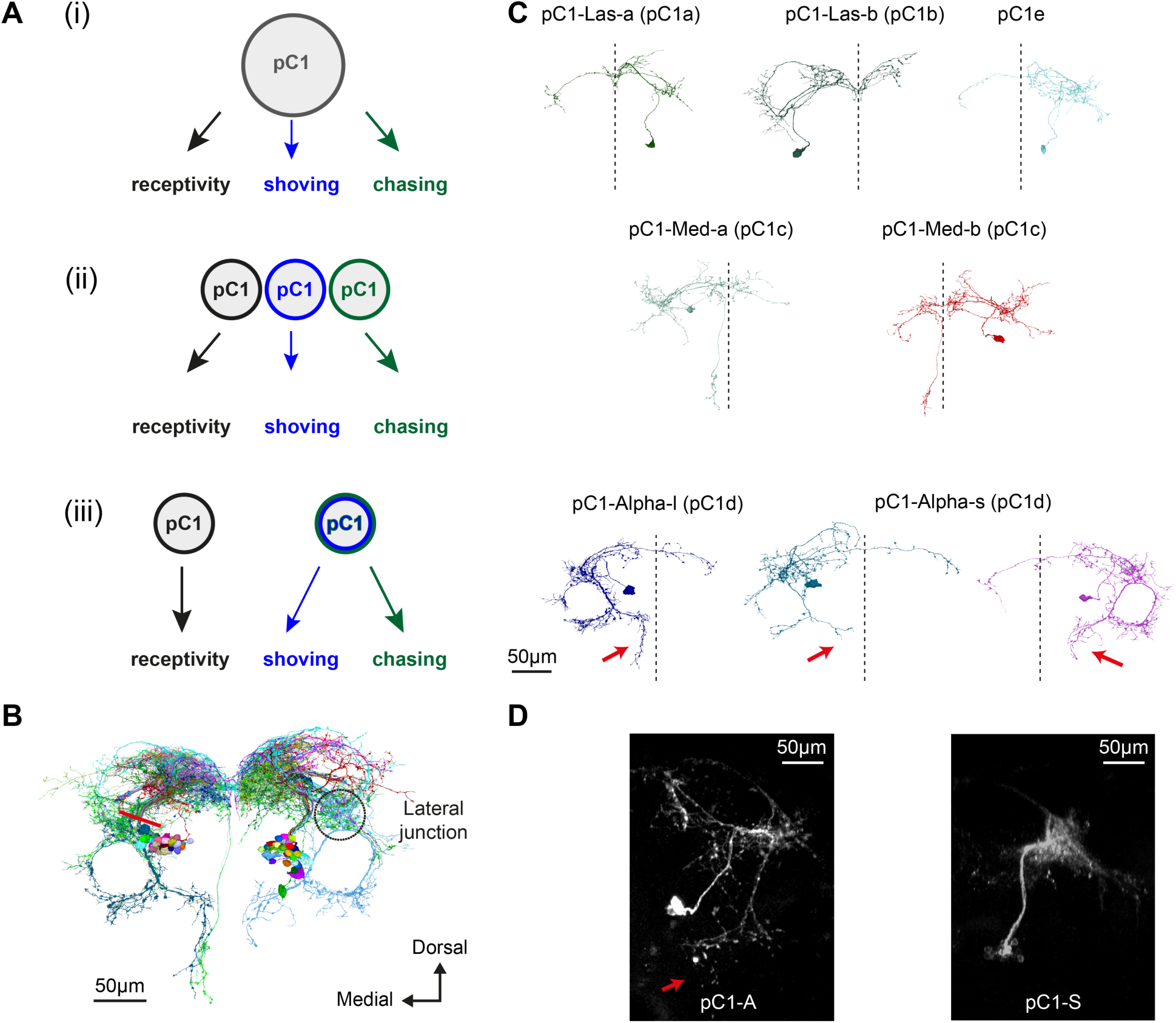
Defining pC1 cell types. **(A)** Three schematic models are shown for the control of pC1-Int neurons over female receptivity/shoving/chasing. In (i), a homogenous pC1 population drives three downstream centers, and in (ii) and (iii) pC1 is a heterogenous group, with different behaviors controlled by different pC1 subsets. **(B)** EM tracing of all neurons that pass through a cross section in the pC1 bundle (red line; see Fig. S3A-B), including neurons that project to the lateral junction (considered pC1 cells), and neurons that do not project to the junction (not considered pC1 cells; see Fig. S3C-D). **(C)** 7 pC1 cell types. pC1-Las-a,b (Las for ‘Lasso’), pC1-Med-a,b (differ in their medial projection; ‘Med’ for medial, they both have a long medial projection), pC1-Alpha-s,l (short and long medial projections (red arrows) and pC1e (named as in Wang et al. 2020). Names in parentheses are the ones used in Wang et al., 2020. **(D)** pC1-A (left; VT25602.AD; VT2064.DBD; n = 7, 2 ± 0 cells per hemisphere (Wu et al., 2019)) pC1-S (right; R71G01.AD; DSX.DBD; n = 8, 7.7 ± 5 cells per hemisphere) neurons expressing GFP and labeled with anti-GFP, see Table 1 for full genotype. pC1-A has a medial projection (red arrow), similar to pC1-Alpha-l neurons found in EM (C); the medial projection was found in 7/7 imaged pC1-A female brains, in both hemispheres. This projection was not found in pC1-S imaged female brains (8/8 brains) - neurons labeled in pC1-S resemble pC1-e neurons found in EM (C).

We systematically checked all cell segments that pass through a cross section in the pC1 bundle (Fig. 2B, red line, Supp Fig. 3A-B) and excluded neurons that do not project to the lateral junction (Supp Fig. S3C-D), as all pC1 cells characterized so far project to the lateral junction (Kimura et al., 2015; Rezával et al., 2016; Wu et al., 2019; Zhou et al., 2014). We sorted pC1 cells manually based on morphology and found 7 major pC1-like cell types in the EM volume (Fig. 3C) - including all 5 pC1 types found from manual tracing in the same EM volume (Wang et al., 2020). Two types have a ring dorsal to the lateral junction (‘pC1-Las-a/b’; Las for Lasso; similar to pC1a,b in Wang et al. 2020), two have a long medial projection (‘pC1-Med-a,b’; Med for Medial; pC1-Med-a is similar to pC1c in Wang et al. 2020), one has dense processes medio-dorsal to the lateral junction (pC1e; same as in Wang et al. 2020). Two other pC1 types have a horizontal projection medial to the ‘ring’ (Yu et al., 2010), similar to pC1 cells previously named ‘pC1-Alpha’ (Wu et al., 2019). We termed these ‘pC1-Alpha-l/s’ for long/short based on the length of the medial vertical projection (Fig. 3C, red arrows; pC1-Alpha-l is similar to pC1d in Wang et al. 2020). The pC1-Int intersection labels ∼7 cells per hemisphere in the brain, and includes the pC1-Alpha-l medial projection; it also has expression in the VNC (Supp Fig. S3E, (Zhou et al., 2014)).

### Different pC1 subtypes affect receptivity versus chasing and shoving

We used genetic intersections to label two non-overlapping pC1 subpopulations: the first labels 2 pC1-Alpha neurons per hemisphere and no cells in the VNC (Wu et al., 2019) - we refer to this line as ‘pC1-A’ (Fig. 3D and Supp Fig. S3F), the second intersection does not label any pC1-Alpha cells (‘pC1-S’, a split-Gal4 intersection between R71G01.AD and DSX.DBD (Pavlou et al., 2016); Fig. 3D and Table 1). The pC1-S line labels 7.7±4 cells per hemisphere in the brain, and no cells in the VNC (Supp Fig. S3G). The pC1-S morphology appears similar to pC1e.

Next, we tested if activation of these two non-overlapping pC1 sub-populations drives persistent behavioral phenotypes. Following activation, neither pC1-A nor pC1-S had an effect on female receptivity (Fig. 4A, D), suggesting that a third population of pC1 cells in the pC1-Int driver line is responsible for driving female receptivity. Activation of pC1-A drove both shoving and chasing (Fig. 4B-C), while activation of pC1-S did not (Fig. 4E-F). This is consistent with model 3 (Fig. 3A). The mean shoving probability following pC1-Int activation (Fig. 4B, yellow diamond) was higher than the mean shoving probability following pC1-A activation (Fig. 4B bars), indicating that pC1 cells other than pC1-Alpha enhance female shoving. On the other hand, female chasing more than doubled following pC1-A activation in the d0 condition (Fig. 4C), compared with pC1-Int activation (Fig. 4C, yellow diamond), suggesting that pC1 cells other than pC1-Alpha suppress female chasing. To determine if the same subset of pC1-Int cells affect both female receptivity and female shoving/chasing, we examined the probability of shoving and chasing relative to receptivity state (Fig. 4G-H). We found that following pC1-Int activation, females that eventually copulated (receptive) showed a higher level of shoving compared with females that did not eventually copulate (unreceptive), though this effect was not statistically significant. In contrast, receptive females showed a reduced level of chasing. This suggests that cells within the pC1-Int line that control receptivity modulate the amount of female shoving and chasing (Fig. 4I).

**Figure 4:**
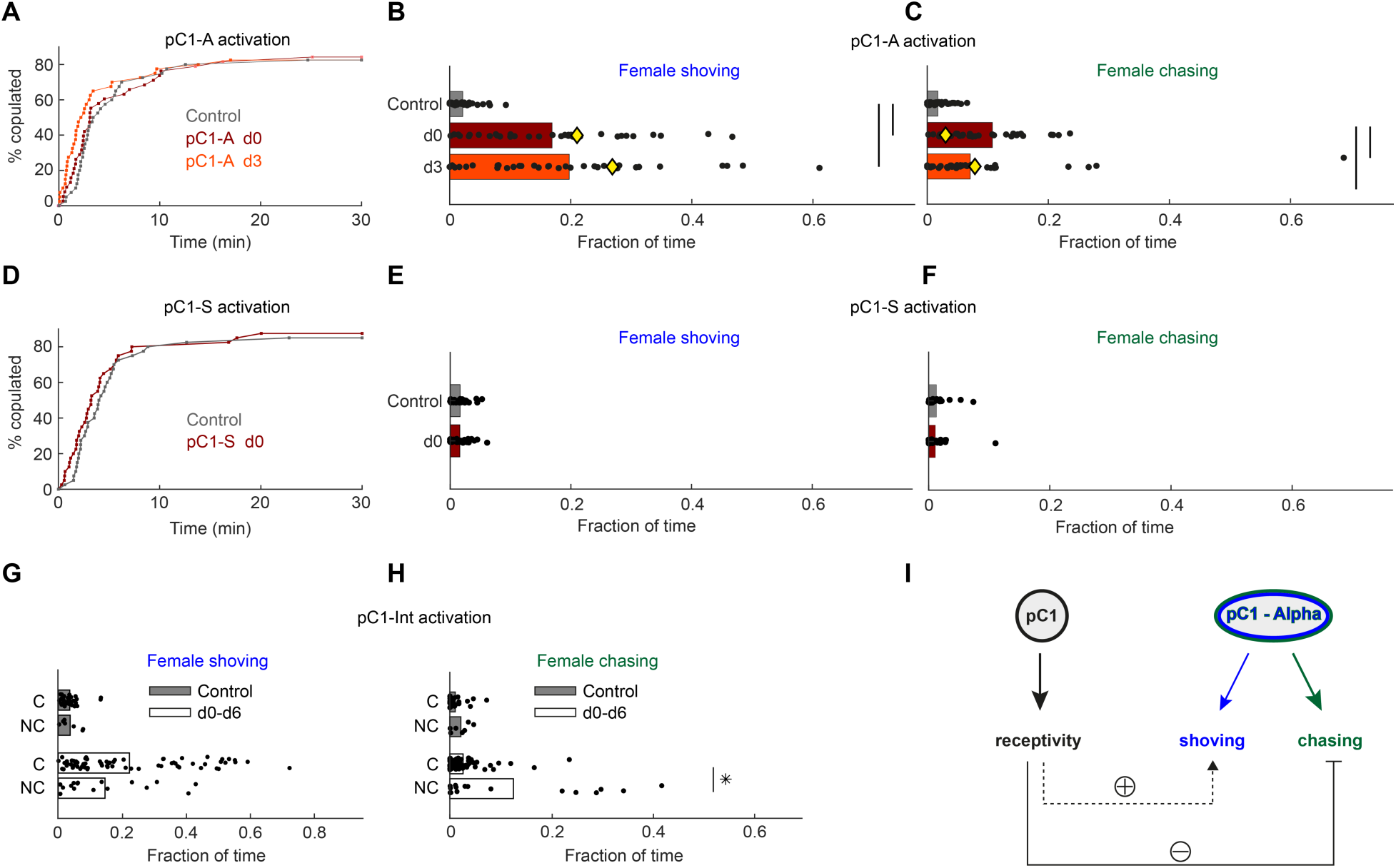
pC1-Alpha neurons drive female shoving and chasing, but do not affect receptivity. **(A)** pC1-A activation did not affect copulation rate in either the d0 or d3 conditions (N = 40, 38, 40 for control/d0/d3; P = 0.79 or 0.29 for control vs d0 or d3; Cox proportional hazards regression, see Methods). **(B, C)** Shoving (B) and chasing (C) probabilities (control/do/d3: 0.02/0.17/0.20 and 0.018/0.11/0.07 for shoving and chasing) were significantly higher in both the d0 and d3 conditions compared to control (two-sample t-test; *P<0.05). Female shoving probability was lower following pC1-A activation, compared with pC1-Int activation (yellow diamond = mean from Fig. 2E). Chasing probability was higher following pC1-Alpha activation in the d0 condition compared with pC1-Int activation (yellow diamond = mean from Fig. 2E). **(D-F)** same as (A-C), but for pC1-S activation. pC1-S did not affect neither copulation rate (D) nor shoving (E) or chasing (F) probabilities (control/do: 0.02/0.02 and 0.01/0.01 for shoving and chasing). **(G)** Fraction of frames with Shoving for copulated (C) and non-copulated (NC) pairs for all experimental conditions taken together (d0-d6). Each dot is a single pair, and the bar value is the mean over all pairs (P = 0.92 and 0.13 for control and d0-d6). **(H)** Same as (G), for female chasing (P = 0.13 and control and P < 10^-5^ for d0-d6). **(I)** A summary schematic. Female pC1-Alpha cells drive persistent shoving and chasing, while not affecting female receptivity. Female receptivity, controlled by a separate (unknown) pC1 subset, suppresses female chasing, while possibly enhancing female shoving.

### Activation of pC1-Alpha neurons drives persistent neural activity

Persistent neural activity is defined as activity that continues after a triggering stimulus goes away (Zylberberg and Strowbridge, 2017). To relate our findings above to persistent neural activity, we activated either pC1-A or pC1-S neurons using the similar pattern of optogenetic stimulation (for 5 minutes, see Methods) that drove persistent changes in behavior (Fig. 1E), and imaged responding cells via GCaMP6s expressed pan-neuronally (Fig. 5A). To compare activity across flies and to map activity onto a reference atlas, we used a recently developed pipeline for two-photon volumetric calcium imaging, motion correction, registration, and region of interest (ROI) segmentation (Pacheco et al., 2019), and scanned the entirety (in the z dimension) of a dorsal portion of a single brain hemisphere in each fly (Fig. 5A). Neuronal activity was measured during the 5 minutes of optogenetic activation in addition to 9.5 minutes following activation offset (Fig. 5B; Supp Movie S4). We found that out of 47,882 ROIs segmented across 28 brains (Fig. 5B; see Methods), 4254 ROIs had significant responses to optogenetic stimulation (Fig. 5C; Ft1 > 3***σ***_0_, ***σ***_0_ = standard deviation of activity during baseline). These ROIs were then clustered based on response patterns (Fig. 5C-D). We found transient responses - ROIs with elevated activity during the optogenetic stimulus (t1), but not following the stimulus (t2) - that could be grouped into two clusters (response types 3 and 4). However, we also found responses with sustained activity lasting at least 5 minutes after the optogenetic stimulus offset (Ft2 > 3***σ***_0_, see Methods). These persistent responses also clustered into two response types (response types 1 and 2). While response type 1 had low spatial consistency across animals, response types 2-4 showed higher spatial consistency, and the spatial distribution of ROIs differed between controls, pC1-S, and pC1-A activated flies (Fig. 5E). pC1-A activation drove persistent activity (response type 2) in more than 30% of the imaged flies, and in 24.7 times more voxels than in controls, and 6.8 times more voxels compared to pC1-S activation (Fig. 5E). The temporal dynamics of persistent neural activity (Fig. 5C), continuing to at least 5 minutes following stimulation, is consistent with our observation of female shoving and chasing of a male introduced 6 minutes after stimulation offset (Fig. 2E-F).

**Figure 5:**
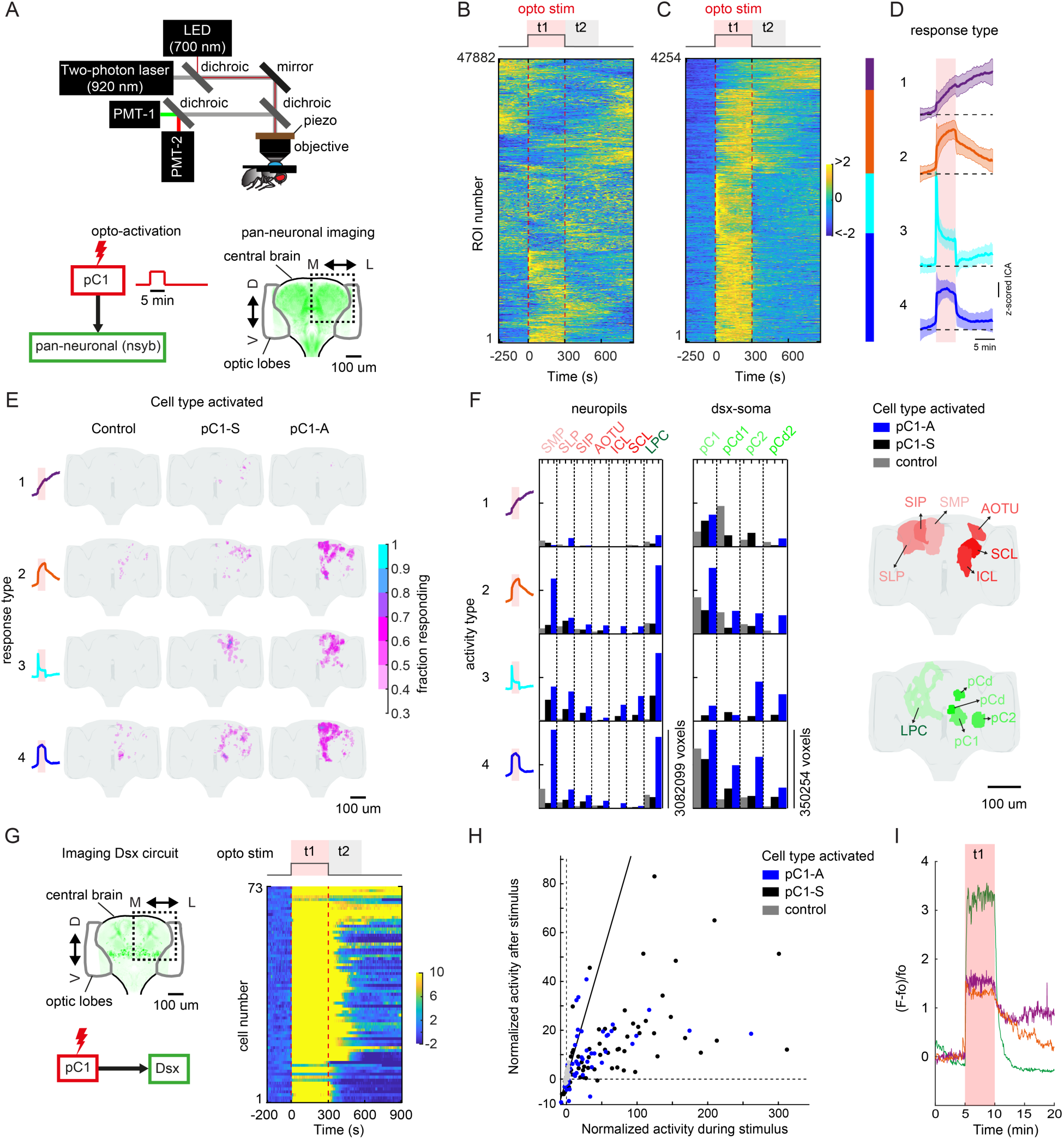
pC1 neurons drive persistent neural activity in the central brain. **(A)** Experimental setup. pC1 cells (pC1-A or pC1-S) expressing csChrimson were activated through the objective using an LED (700nm). GCaMP6s and TdTomato were expressed pan-neuronaly using the nsyb driver, and a custom-designed two-photon microscope was used to image brain activity before, during and after pC1 activation (see Table 1 for genotypes and Methods for more details on the experimental setup). **(B)** Brain activity recorded in response to optogenetic stimuli (N = 28 flies). GCaMP6s signal was motion corrected and 3D-ROI segmented based on correlated activity in neighbor voxels (see Methods). The z-scored signal of all ROIs (N = 47882 ROIs from both pC1-S and pC1-A activation and control experiments) are color coded (in units of standard deviations, see scale bar), and shown 5 minutes before activation, during activation (t1), and 9.5 minutes post-activation (t2 marks the first 5 minutes post-activation). ROIs are sorted based on hierarchical clustering of the temporal dynamics. Red dashed line depicts the optogenetic stimulus onset and offset. **(C)** pC1 activation evokes transient and persistent activity. ROIs were sorted based on mean z-scored activity during (t1) and after photoactivation (t2). We found 4254 responsive ROIs, defined as ROIs with Ft1 > 3***σ***_o_ (***σ***_o_ - standard deviation during baseline, Ft1 is the mean fluorescence during t1). These ROIs were split into transient (Ft2 ≤ 3***σ***_o_, blue and cyan; Ft2 is the mean fluorescence during t2) or persistent response types (Ft2 > 3***σ***_o_, red and purple). Lines within each response type were sorted based on hierarchical clustering of temporal dynamics. The 4254 ROIs were sorted to Ft2 ≥ 3***σ***_o_ (red and purple) or Ft2 < 3***σ***_o_ (blue and cyan). These response types were further clustered into response types 1-4 (red, purple, blue, and cyan) based on temporal dynamics (see Methods). **(D)** Mean ± SD for response types 1-4. In response types 1 and 2 the activity level (calcium response) persists after activation offset, while for types 3 and 4, the activity is high during, but not after photoactivation. **(E)** Maps of transient and persistent activity types. ROIs from response types 1-4 per animal were registered to an in vivo intersex atlas (Pacheco et al., 2019) to generate probability density maps across animals per brain voxel (each voxel is 0.75×0.75×1 µm^3^). Maps of activity are overlaid onto the brain template, color coded by the fraction of flies showing activity on each voxel (ranging from 30-100%). We considered a voxel to consistently have a particular response type if active in over 30% of flies. Response type 2 shows persistent activity following pC1-A activation, and occupies 4.3% of a single hemisphere following pC1-A activation compared to 0.6% following pC1-S and 0.2% in control flies. **(F)** Brain regions containing transient and persistent response types. We used both standard anatomical segmentation of the in vivo brain atlas and segmentation of Dsx circuit (into LPC neuropil - Lateral Protocerebral Complex (Yu et al., 2010) - and major groups of cell bodies (pC1, pC2, pCd1, pCd2)) to assign ROIs to neuropils or Dsx related domains (see spatial locations in green on the right). For each brain region, we calculated the average number of voxels or volume (across-individuals) occupied by all ROIs belonging to each response type for each condition (pC1-A/pC1-S activation, or controls). Neuropils were sorted by the number of voxels with response type 2 following pC1-A activation, and the top 6 neuropils (names in red, see spatial location in red on the right) are shown. pC2m and pC2l are shown together as pC2, as they are not always spatially separable in females. **(G) Left:** Experimental design of pC1 activation and readout of activity in Dsx cell bodies. **Right:** Normalized activity of pC1 cell bodies ((F(t) - F0)/***σ***_o_; F0 and F(t) are mean Fluorescence during baseline and fluorescence over time, respectively). Chrimson and TdTomato are expressed in pC1-A or pC1-S cells. GCaMP6s is expressed in Dsx+ cells. We show 73 traces with high correlation between the stimulus pattern and Calcium response (>0.5, see Fig. S4C). **(H)** Mean Calcium response during t1 (x-axis) versus during t2 (y-axis) are shown for all conditions. Normalized activity is defined as (F - Fo)/***σ***_o_, where Fo is the mean activity during baseline, ***σ***_o_ is the standard deviation during baseline, and F is the mean activity during t1 for x and t2 for y. Each dot represents a single cell. All recorded pC1 cells are shown. **(I)** Example traces of (F - Fo)/Fo from 3 individual pC1 cells, showing different profiles of Calcium decay profile after stimulus offset.

Making use of anatomical segmentation of an *in vivo* brain atlas to which all ROIs were registered (Pacheco et al., 2019), we evaluated the distribution of pC1-elicited activity by brain neuropil (Ito et al., 2014). Persistent activity was clustered in the posterior-dorsal portion of the brain spanning the Superior Medial, Lateral and Intermediate Protocerebrum (SMP, SLP, SIP), the Anterior Optic Tubercle (AOTU), and the Inferior and Superior Clamp (ICL and SCL; see Fig. 5F and Supp Fig. S4A); these brain regions contain a large number of projections from sexually dimorphic neurons expressing either Doublesex or Fruitless (Rideout et al., 2010; Yu et al., 2010). In order to measure the overlap of pC1-elicited activity with Dsx+ neurons, we generated anatomical labels for the lateral protocerebral complex (LPC), a diffuse brain area to which all Dsx+ neurons send their projections, and also for all major groups of Dsx+ somas (pC1, pC2, pCd1, and pCd2) within the *in vivo* brain atlas (Fig. 5F, see Methods). We found that ROIs with persistent activity (response type 2) overlap with the LPC, in addition to the regions occupied by pC1 somas, and to a lesser extent with regions occupied by pC2, pCd1 and pCd2 somas, suggesting that Dsx+ neurons carry persistent activity.

To directly test this, we expressed GCaMP6s only in Dsx+ neurons (Fig. 5G). We activated either pC1-A or pC1-S for 5 minutes (Fig. 5G and Supp Movie S5) and recorded activity in 619 cells (we manually identified somas) across 42 flies. We examined the responses during (t1) and after (t2) optogenetic stimulation (same as for the pan-neuronal dataset), and compared these responses to controls in which pC1 neurons were not activated (N = 10 flies, 192 ROIs) (Fig. 5H). A number of pC1 cells showed strong persistent activity (Fig. 5G; same definition as for the pan-neuronal screening, Ft2 > 3***σ***_0_) following optogenetic activation of either pC1-A or pC1-S neurons, although there was some heterogeneity in responses across the pC1 cells (Fig. 5G-I). We did not observe persistent activity in any other Dsx-expressing cell types (Supp Fig. S4B-D), including the pCd1 cells, previously shown to be necessary for P1 induced persistent activity in males (Jung et al., 2020; Zhang et al., 2019).

In sum, our pan-neuronal imaging reveals a large difference in the numbers and spatial distribution of ROIs following pC1-S versus pC1-A activation (Fig. 5E) - this is consistent with our behavioral experiments in which we observed that pC1-S and pC1-A activation had different effects on female behavior. However, imaging of individual Dsx-expressing neurons (Fig. 5G-I) showed that similar numbers of pC1 cells contain persistent activity following activation of either pC1-S or pC1-A neurons. This could be because different subsets of pC1 neurons are active following pC1-S versus pC1-A activation, and these different neurons have different contributions to behavior. Nonetheless, our imaging experiments reveal pC1-A driven persistent neural activity lasting for minutes following optogenetic activation in specific cells and brain regions.

### pC1-Alpha is reciprocally connected to a specific subset of Fruitless+ neurons

Recurrent connectivity between neurons is known to support persistent neural activity (Goldman-Rakic 1995; Zylberberg and Strowbridge 2017; Major and Tank 2004). We searched for recurrent connections between pC1-Alpha and other neurons in the female brain, using automated reconstruction of all neurons within an EM volume of an entire adult female brain ((Zheng et al., 2018); flywire.ai, manuscript in preparation). This volume contains both brain hemispheres, enabling complete reconstruction of single pC1-Alpha cells that send projections across the midline (Fig. 3C). We focused on a single pC1-Alpha-long cell (Fig. 6A) that has a long vertical medial projection (see also (Wu et al., 2019). We manually detected all synaptic connections, as done previously (Felsenberg et al., 2018; Sayin et al., 2019; Zheng et al., 2018). After excluding very weak connections (using 3 synapses as a threshold, see Methods), we counted 417 presynaptic and 421 postsynaptic sites (Fig. 6B and Supp Movie S6). We found mostly output synapses in the dorsal projection that crosses the midline, mostly input synapses in the medial projection and on the medial side of the ring, and mixed input/output synapses in the rest of the ring, including in lateral junction (Fig. 6B). This finding is generally in agreement with previous mapping of inputs/outputs of pC1 neurons based on molecular tags (Kimura et al., 2008; Zhou et al., 2014). We sorted all pC1-Alpha synaptic partners by cell type, based on morphology, and examined the distribution of synapses by type for input (presynaptic partners) and output (postsynaptic partners) neurons separately (Fig. 6C). We found a difference between the input and output distributions: while 3 cell types encompass almost half of the output synapses (49.4%), the 3 most common input cell types encompass only 30.5% of all input synapses (Fig. 6C-D, Supp Fig. S5A). This asymmetry may suggest that pC1-Alpha neurons act as a convergence node of multiple inputs that drive a smaller number of outputs.

**Figure 6:**
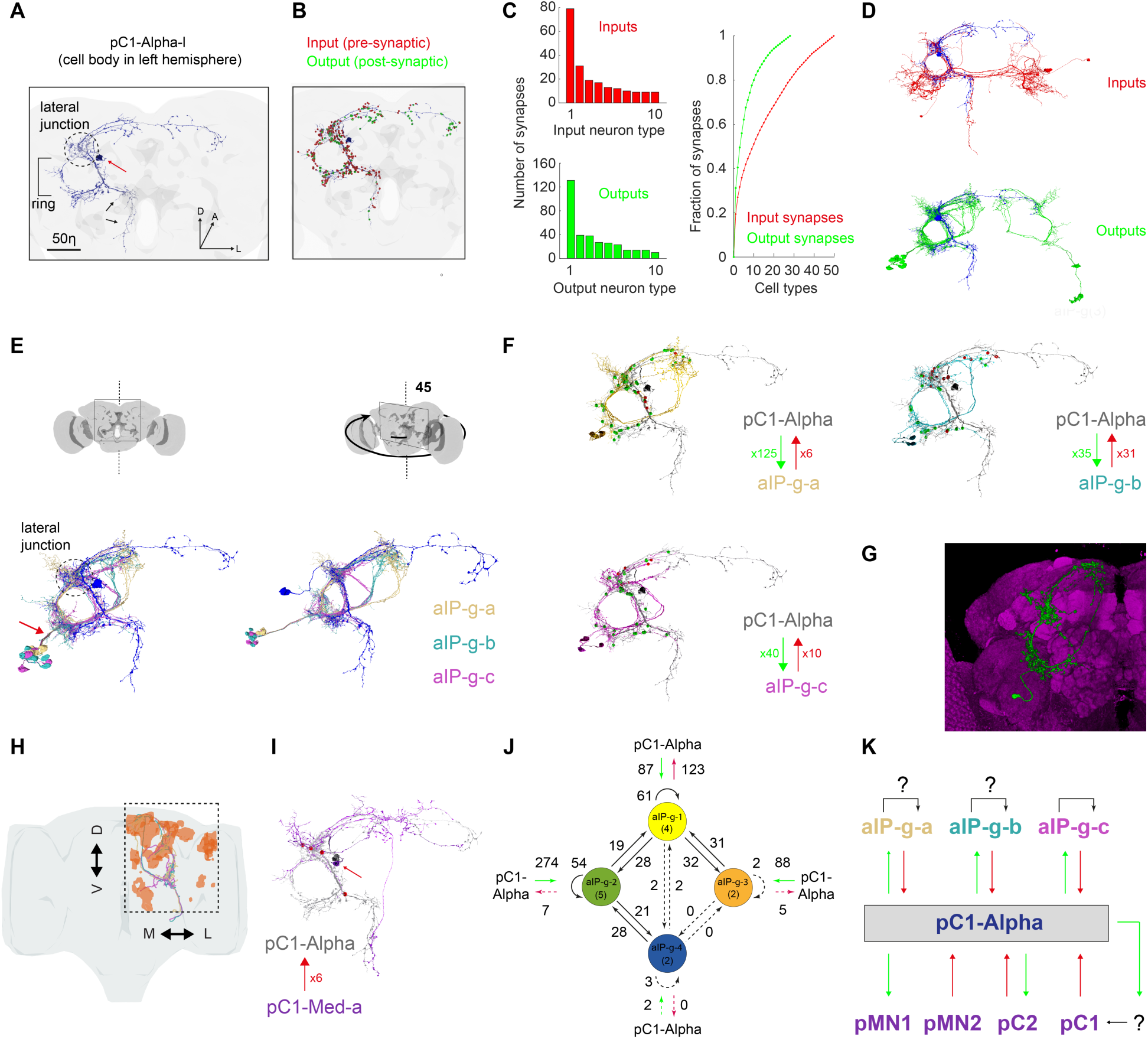
The connectome of pC1-Alpha reveals recurrent connections with aIP-g neurons. **(A)** pC1 Alpha-l cell from FAFB volume after automatic segmentation and manual proofreading. The cell body is marked with a red arrow, and the pC1-Alpha medial projection (that does not exist in other pC1 types) is marked with black arrows. **(B)** Same cell as in (A), with manually detected synapses. Presynaptic terminals (inputs to pC1-Alpha-l) are marked in red, post-synaptic terminals (outputs) in green. After excluding segments that are connected with pC1-Alpha-l with less than 3 synapses, we end up counting 417/421 input/output synapses (see also Supp Movie S6). **(C) Left:** pC1-Alpha-l inputs (66 cells) and outputs (50 cells) were classified manually to cell types based on morphology. The number of input (top) or output (bottom) synapses are shown for each type, sorted (separately for inputs and outputs) based on the total number of synapses with pC1-Alpha-l for each type. **Right:** The cumulative fraction of synapses counted as a function of the number of types included (calculated separately for inputs/outputs). The three most common output types encompass 49.4% of the output synapses, while the 3 most common input types encompass 30.5% of all input synapses. **(D)** pC1-Alpha-l major inputs (top) and outputs (bottom). Only the cells that belong to the most common cell types (50% input or output synapses) are shown. One cell is shown per cell type. Note that pC1-Alpha-l has postsynaptic connections with both left (ipsilateral to cells body) and right (contralateral) aIP-g cells. **(E) Left:** Posterior view (same as in (A)) of pC1-Alpha-l (blue) and example aIP-g cells (3/6/5 for types a/b/c - aIP-g neurons were identified in (Cachero et al., 2010), but we define the three subtypes based on projections). **Right:** view rotated by 45 deg, showing the separation between three subtypes of aIP-g cells, that we named aIP-g-a, aIP-g-b and aIP-g-c. **(F)** pC1-Alpha-l (grey) aIP-g-a,b,c cells (same color code as in (E)). pC1-Alpha Input/output synapses with aIP-g are shown in red/green for each type, and the total number of input/output synapses are shown for each group. **(G)** A single clone of a Fru+ neuron from FlyCircuit (http://www.flycircuit.tw/) was found by NBLASTing (Costa et al., 2016) a single aIP-g cell. Cell name is fru-F-200105, VirtualFlyBrain ID VFB_00004510. **(H)** Example aIP-g-a,b,c cells (one of each) using only the automated segmentation (before proofreading), together with the map of regions with persistent activity following pC1-A activation (reproduced from Fig. 5E, response type 2) (brown). Dashed box defines the area imaged (same as Fig. 5A). Overlap between each example aIP-g and Calcium activity was calculated by first voxelizing points along the cell, then looking for the response type (1-4) in each voxel. We found that 30.7/15.5/10.1% of aIP-g-a/b/c overlapped with response type 2 (response type 2 represents 4.3% of the entire central brain volume). **(I)** An example pC1 cell (type pC1-Med-a) that is presynaptic to pC1-Alpha-l (6 synapses were manually detected). **(J)** The number of synapses within and between groups of aIP-g cells based on the fly hemibrain connectome (Xu et al., 2020). The number in parentheses indicates the number of cells per group (aIP-g-1-4). Round arrows indicate within-group connections (e.g., 61 synaptic connections between pairs of aIP-g-1 cells). Dotted arrows are shown for weak connections (under 5 synapses). **(K)** A summary of pC1-Alpha connections with aIP-g and other Dsx+ neurons. As pC1 cells show persistent activity following pC1-A activation, we propose indirect connectivity between pC1-Alpha and other pC1 cell types. aIP-g-c cells are reciprocally connected as shown in Fig. 6J. Potential connections within aIP-g-b or aIP-g-c groups are indicated as a question mark. Connections between neurons belonging to different aIP-g groups are not indicated for simplicity.

The three output types that have the largest number of synapses with pC1-Alpha, share a common morphology (Fig. 6E, Supp Fig. S5B and Supp Movie S7). All three types have projections in the ‘ring’ (Fig. 6A; (Yu et al., 2010)), where they overlap with pC1-Alpha neurons, including dense projections to the lateral junction (Fig. 6E). We found that some neurons from each one of the three types, are also presynaptic to pC1-Alpha, and thereby form a recurrent connection with pC1-Alpha (Fig. 4F). We used NBLAST (Costa et al., 2016; Manton et al., 2019) to search for matches between the EM tracing of neurons in these three neuron types and FlyCircuit neurons ((Chiang et al., 2011)(Osumi-Sutherland et al., 2014); see Methods). The top matches for all three types were Fru+ neurons called aIP-g (Fig. 6G and Supp Fig. S5C-D). This cell type was previously described in males, and its morphology is sexually dimorphic (Cachero et al., 2010). Notably, each one of the three types we used as a seed for a search, resulted in a different aIP-g cell: aIP-g-a, aIP-g-b and aIP-g-c (Fig. 6F, Supp Fig. S5C and Supp Movie S7; we found a total of 6/30/8 aIP-g-a/b/c cells in the left hemisphere), each of which has reciprocal connections with pC1-Alpha. The input and output synapses are asymmetrically distributed - for example, pC1-Alpha has ∼20 times as many output synapses with aIP-g-a as input synapses (Fig. 6F). Above, we overlaid the map of persistent neural activity driven by activation of pC1-A neurons with markers for Dsx+ expression (Fig. 5E-F); from this, we observed that Dsx+ pC1 cells carried persistent activity, and confirmed this result by imaging responses in only Dsx+ cells (FIg. 5G-H). We used a similar strategy to examine the overlap between the persistent activity following pC1-A activation with the projections of individual aIP-g neurons, all registered into the same reference brain (Fig. 6H; see Methods). While persistent activity (response type 2; Fig. 5D-E) spans only 4.3% of the entire brain (Fig. 5E; see Methods), we found response type 2 activity in 30.7/15.5/10.1% of the voxels that include aIP-g-a/b/c example skeletons (Fig. 6H).

Next, we looked at the connectivity between pC1-Alpha and other Dsx+ neurons (including pC1, pCd, pC2, pNM1, and pNM2 (Fig. 1A)), following our observation that pC1-A activation drives persistent activity in other pC1 neurons (Fig. 5F-I, Supp Movie S5), and transient responses in some pCd and pC2 neurons (Supp Fig. S4B-D). We divided pC2-like neurons into 3 different cell types (Supp Fig. S5E), and compared each type to single clones of Dsx+ pC2 neurons (Deutsch et al., 2019) - this analysis suggested that pC2-DR (DR for Double Ring) is likely Dsx+. We did not observe recurrent connections between pC1-Alpha and any pC1 or pC2 like neuron (meeting our criteria for connections; see Methods), indicating that aIP-g neurons may be special in their recurrent connectivity with pC1-Alpha. We did however find non-reciprocal connections between pC1-Alpha and other Dsx+ cell types (Fig. 6I, Supp Fig. S5E-F and Supp Movie S8).

Lastly, we looked at the synaptic connectivity between pC1-Alpha and aIP-g cells in a second EM database that consists of a portion of the adult female brain (the ‘hemibrain’; (Xu et al., 2020)), and found a set of 13 neurons identified as aIP-g ((Xu et al., 2020); in this dataset a single pC1-Alpha has been traced, and is named pC1d (Wang et al. 2020)). These aIP-g cells (denoted as types aIP-g-1-4 in the hemibrain) all share the aIP-g-c morphology (Supp Fig. S5G), and 12 of them are synaptically connected to pC1d (excluding 1 connection with less than 3 synapses). The ratio between the total number of pC1d-->aIP-g and aIP-g-->pC1d synapses is 3.3:1 in the hemibrain, compared to 4:1 (for aIP-g-c) in FlyWire (FAFB), although the absolute number of synapses is significantly higher in the hemibrain, possibly due to the conservative definition we used for manual synapse detection in FAFB (see Methods). We also examined synaptic connections between pairs of aIP-g cells in the hemibrain and found additional recurrent connectivity within the aIP-g group (Fig. 6J). These results indicate that pC1-Alpha serves as a hub within the central brain linking Dsx+ neurons to Fru+ neurons (Fig. 6K). As we observed persistent neural activity in several pC1 cells following pC1-A activation (Fig. 5G-H), we hypothesize the existence of indirect connections between pC1-Alpha and other pC1 cells.

## Discussion

We find that pC1 neurons drive a persistent internal state in the *Drosophila* female brain that modulates multiple behaviors over timescales of minutes (receptivity, responses to male courtship song, aggressive behaviors, and male-like courtship behaviors (Figs. 1-2 and 4)). The behavioral effects we observe are in line with the effects of ‘emotion states’ observed in other animals, such as mice, fish, and primates (Anderson and Adolphs, 2014; Kunwar et al., 2015; Posner et al., 2005; Russell, 2003; Woods et al., 2014). In general, effects on behavior of such emotion states are thought to scale with levels of persistent neural activity (Kunwar et al., 2015; Lee et al., 2014) - this could be influenced either by the level of stimulation that drives the persistent state or the amount of time that lapses following stimulation. Several studies have explored the former (varying the level of stimulation) and have found in both mice and flies, that weak activation drives different persistent behaviors versus strong activation (Hoopfer et al. 2015; Lee et al. 2014). Here, we explore the latter (varying the delay after stimulation) - this was done to separate the levels of persistent activity (which decay during the delay period, as we observe in our neural imaging experiments (Fig. 5)) from stimulus-driven behaviors (in our case, behaviors produced towards a male fly). We found that the duration of the delay between pC1 neuron activation and introduction of a male fly affects the amount and type of behaviors that are produced.

Our study also provides new insight into the neural mechanisms that contribute to changes in state on timescales of minutes (Figs. 5 and 6). We used pan-neuronal imaging with registration to map responses that continue following pC1 optogenetic activation (previously this technique had only been used to map sensory activity (Pacheco et al., 2019) and spontaneous activity (Mann et al., 2017)). We found that activation of pC1-Alpha neurons drives robust persistent neural activity throughout the posterior dorsal regions of the central brain (known to contain the processes of sexually dimorphic neurons (Cachero et al., 2010; Kimura et al., 2015)), lasting for minutes following activation. This is consistent with our behavioral observations - females still show elevated shoving and chasing even following a 6 minute delay between optogenetic activation and the introduction of a male fly. Importantly, whether or not pC1 neurons themselves carry persistent neural activity has been debated (Inagaki et al., 2014; Jung et al., 2020; Zhang et al., 2018). Here we find that in females, pC1 neurons do indeed carry persistent neural activity in response to our activation protocol (Fig. 5H). Finally, by mapping all synaptic partners of a pC1-Alpha neuron, we find a recurrent circuit motif that may underlie the persistent neural activity.

### pC1 neurons drive both aggression and receptivity in *Drosophila* females

We used unsupervised methods to identify the most prominent behaviors (beyond receptivity and responses to courtship song (Fig. 1)) produced by activating pC1 neurons in virgin females - these include behaviors that resemble male courtship (female chasing the male) and aggression (female shoving the male) (Fig. 2). Both behaviors are not typically observed in mature virgin females interacting with a male (see controls in Fig. 2); this suggests that sensory cues from the virgin male do not inhibit these aberrant behaviors, but rather may enhance the effects of pC1 activation (Fig. 2F). Activation of a subset of pC1 neurons is also known to drive aggressive behaviors towards females during stimulation (Palavicino-Maggio et al., 2019), but whether the quality of aggression generated towards males versus females (and any persistence) is similar remains to be determined. As one of our manually scored behaviors, ‘female approaching’ (Fig. S2C), begins from a distance greater than 4 body lengths from the male fly (a distance at which it may be difficult to discern male from female (Borst, 2009)) and often ends with shoving or circling (see Supp Movie S3), we hypothesize that pC1 activation most likely drives persistent behaviors towards another fly, and not specifically a male or female fly. Our approach for quantifying behaviors should facilitate further dissection of the sensory cues that promote persistent changes in behavior.

What is the role of female aggression? Female aggression, whether towards males or females, has been previously reported across model systems (Huhman et al., 2003; Stockley and Bro-Jørgensen, 2011; Woodley et al., 2000). In *Drosophila*, female-female fights over food source are strongly stimulated by the receipt of sperm at mating (Bath et al., 2017), and include both patterns that are common with male aggression (such as shoving and fencing) and female-only patterns (Nilsen et al., 2004). The behavioral changes in our study do not mimic those in a mated female, as we also observe that pC1 activation drives enhanced receptivity. While we do not yet know which pC1 cell types control receptivity, our work reveals a separation: pC1-Alpha neurons are sufficient to drive shoving/chasing, but do not affect receptivity (Fig. 4A-C), while separate pC1 neurons that control receptivity modulate the pathways that control chasing and aggression (Fig. 4G-H). Interestingly, previous work in male flies suggests a separation in pC1 subsets that control courtship versus aggression (Koganezawa et al., 2016), with reciprocal inhibitory influences between persistent courtship and aggression, following pC1 activation (Hoopfer et al., 2015). Though the phenotypes are sex-specific (male singing vs female receptivity; male tussling vs female shoving), and the pC1 subsets driving these behaviors are sex-specific (P1 in males, pC1-Alpha in females), this suggests some common architecture. Ultimately, comparing the connectomes of male and female brains, combined with functional studies, should elucidate both similarities and differences.

### Recurrent circuitry and persistent neural activity

FlyWire enabled a systematic search for all of the synaptic partners of a single pC1-Alpha cell, and revealed reciprocal connectivity with aIP-g cells. We then identified aIP-g cells in the hemibrain (Supp Fig. S5G) and used this resource to find connections between pairs of neurons within this group (Fig. 6J). Because FlyWire is based on the FAFB dataset of the entire adult female brain (Zheng et al., 2018), our search for synaptic partners of the pC1-Alpha cell could completely cover both hemispheres. Because the hemibrain contains automatically detected synapses (Xu et al., 2020), we were able to rapidly find connections between aIP-g cells. Adding automatically detected synapses to FlyWire will be straightforward, especially given that such data has recently been released for the FAFB dataset (Buhmann et al., 2019).

Our identification of a strongly recurrent circuit between pC1-Alpha and aIP-g neurons is in line with prior work connecting recurrent circuits to persistent activity. For example, recurrent networks in rats (Chen et al., 2017), fish (Aksay et al., 2007), and flies (Cognigni et al., 2018; Jung et al., 2020; Kim et al., 2017; Zhang et al., 2019; Zhao et al., 2018) contribute to persistent activity underlying short-term memory, although the precise mechanism, whether relying on intrinsic properties of individual neurons or on the connectivity within neural circuits, may vary across systems (Barak and Tsodyks, 2007; Major and Tank, 2004; Zylberberg and Strowbridge, 2017). While persistent activity was shown to outlast stimulation for minutes in slices or in isolated brains (e.g., (Egorov et al., 2002; Lyutova et al., 2019)), short-term or working memory related persistent activity *in vivo* typically lasts for seconds (Aksay et al., 2007), even when the delay period between stimulus presentation and behavior is variable (Park et al., 2019). Internal states underlying social behaviors, as we have shown here, persist on very long timescales of many minutes. Mechanisms to support such long timescales have been proposed to be hormonal (McEwen et al., 2015; Sapolsky et al., 2000) or peptidergic (Bargmann, 2012; Marder, 2012), but recent work in male mice (Kennedy et al.) and male flies (Jung et al., 2020) has additionally proposed the role of recurrent circuitry as the underlying mechanism. Here, we confirm this by showing that a single cell type, pC1-Alpha, in the *Drosophila* female brain has both strong recurrent connections and drives persistent neural activity lasting for minutes following stimulation. While we don’t yet know if the aIP-g neurons are required to maintain persistent neural activity, this study provides an entry point for studying such mechanisms.

In sum, our work builds on prior studies showing that *Drosophila* females play an active role in courtship (Coen et al. 2014). Here, we find a novel arousal state in females, which, similar to males, is driven by sex-specific neurons and mediated by minutes-long persistent neural activity. By leveraging new resources for circuit mapping in female brains, we highlight opportunities for linking neural function and behavioral states in this model system.

## Author Contributions

DD and MM designed the study. DD, LE-R, RF, and EI collected and analyzed behavioral data. DD and DAP collected and analyzed neural imaging data. DD, LE-R, and ATB proofread neurons and identified synaptic connections in FlyWire. TP and AC wrote software for tracking of fly orientation and movement. CM, SD, TM, RL, KL, NK, DI, MC, AK, CJ, WS, JW, and HSS developed FlyWire with help from DD, LE-R, and MM. DD, LE-R, and DAP made figures. DD and MM wrote the manuscript, with feedback from all authors.

## Acknowledgements

We thank Barry Dickson, Annegret Falkner and Christa Baker for comments on the manuscript and the entire Murthy lab for helpful discussions. We thank Barry Dickson for sharing the pC1-A split GAL4 line ahead of publication. We thank Stephan Thiberge for assistance with two-photon imaging, Nat Tabris for assistance with software development, and Shruthi Ravindranath for assistance with identifying neurons in FlyWire. This study was supported by an NIH BRAIN Initiative RF1 MH117815-01 to MM and HSS and an NIH BRAIN R01 NS104899 and HHMI Faculty Scholar award to MM.

## Methods

### Fly stocks

All flies were raised at 25°C on standard medium in a 12 hr light/12 hr dark cycle at 60% relative humidity. Female flies used for optogenetic experiments were fed with food that contained all-trans-retinal (Sigma R2500-100MG; ATR concentration is 1 mM) for a minimum of three days post eclosion. Control flies were raised on regular fly food after eclosion. Both experimental and control female flies used for optogenetic experiments, were reared post-eclosion in dark blue acrylic boxes (acrylic available from McMaster-Carr, #8505K92).

A list of full genotypes for the flies used in each Figure can be found in Supplementary Table 1. Fly strains used in this study include: wild type NM91 (Coen et al., 2014) from the Andolfatto group at Columbia University, Dsx-GAL4 (Rideout et al., 2010) and Dsx-Gal4.DBD (Pavlou et al., 2016) from Stephen Goodwin, R71G01-p65.AD;MKRS/TM6B,tb (#70798; (Dionne et al., 2018)) and GMR57C10-LexA (#52817) from Gerry Rubin, 10xUAS-Syn21-Chrimson-tdTomato 3.1 [attP18], 13xLexAop2-IVS-Syn21-opGCaMP6s from Allan M. Wong (Hoopfer et al., 2015). w+,NorpA[36],20xUAS-csChrimson-mVenus[attp18];CyO/Sp;MKRS/TM6B,tb is from Vivek Jayaraman, VT25602.p65ADZp; VT2064.ZpGAL4DBD (Wu et al., 2019) and UAS>STOP>TNT (Stockinger et al., 2005) from Barry Dickson, R41A01-LexA (Zhou et al., 2014), Dsx-LexA::P65 (Zhou et al., 2015) from Bruce Baker, 8xLexAop-mCD8tdTomato from Yuh Nung Jan, and UAS(FRT.mCherry)ReachR [attp5] (now Bloomington #53743) from David Anderson (Inagaki et al., 2014). The following flies came from the Bloomington *Drosophila* stock center: UAS-2xEGFP;Dsx-Gal4 (#6874), R71G01-LexA::p65 [attp40] (Pan et al., 2012) (#54733), w[*]; P{UAS(FRT.w[+mW.hs])TeTxLC}10/CyO (#28842; Keller et al. 2002), 8xLexAop-FLP [attp2] (#55819) and 13xLexAop-GCaMP6s (#44590) (Pfeiffer et al., 2010).

### Behavioral experiments

All behavioral experiments were carried out using a behavioral chamber (diameter ∼25mm) tiled with 16 microphones (NR23158, Knowles Electronics; Fig. 1B and Supp Movies 1-3), and connected to a custom amplifier (Arthur et al., 2013)). Audio signals were recorded at 10KHz, and the fly song was segmented as previously described (Arthur et al., 2013; Coen et al., 2014). A point grey camera (FL3-U3-13Y3M; 1280X960) was used to record fly behavior from a top view (see Figs. 1B, S1A) at 60 frames per second using custom written software in Python and saved as compressed videos (H.264). Virgin females (see Table 1 for genotype used in each experiment) and wild type NM91 virgin males (both males and females were 3-7 days old) were used for all behavioral experiments. For inactivation experiments (Figs. 1C-D, S1C), a male and female were introduced into the behavioral chamber simultaneously. For activation experiments, a female was positioned in the behavioral chamber, and red light (a ring of 6 LEDs, 627nm, LuxeonStar) was then delivered at 1.1mw/mm^2^ (±5% across the chamber) at 100Hz (50% duty cycle) for 5 minutes (Fig. 1E). Following stimulus offset, a male was introduced with either no delay (d0), or after a 3- or 6-minute delay (d3, d6). We collected data from the time the male was introduced (t = 0) until 30 minutes or when the flies copulated, whichever came first. We choose to use ReachR (Inagaki et al., 2014) for activation experiments rather than using the more sensitive red shifted channelrhodopsin csChrimson (Klapoetke et al., 2014) to minimize optogenetic activation due to background light. The percent of flies that copulated as a function of time (Fig. 1C,F, Fig. S1D) was calculated from the time the male was introduced (t = 0) and until t = 30 minutes.

### Tracking centroids and headings

Each video frame was analyzed by first finding the fly centroid, then detecting the body parts and extracting heading for each fly. Having microphones at the chamber bottom results in a highly inhomogeneous background (Fig. S1A) posing a major challenge for accurate tracking. The centroid was found in one of two ways that yielded similar results. In the first method, the inhomogeneous background of the video was found by taking the average across all frames. Because the animals move throughout the video, finding the median pixel usually does not contain any pixels containing an animal’s body. However, as animals occasionally sit for long periods, they can become part of the background. To avoid this, we divided the video into 10 shorter videos of equal length and found the median ‘frame’ (median set of pixel values) for each sub-video. We then created a median frame by computing the median across these medians. Each video frame then had this background subtracted to identify pixels that were potentially part of each fly. These pixels were smoothed by a series of operations using the OpenCV Python package and then thresholded. Using OpenCV, we identified all contours surrounding collections of pixels and any smaller or larger than some predefined threshold (less than half the size of a typical ‘fly’ or more than twice its size) were discarded. The remaining pixels were then clustered via k-means. The number of clusters were iteratively increased until the compactness of each cluster reached some threshold. The least-compact clusters were discarded, and the remaining pixels were clustered again with k-means with k=2 to identify the two clusters corresponding to the animals. These clusters were then fit with an ellipse to identify the centroid of each animal. In the second method, we trained a deep convolutional network to detect all instances of individual body parts (head, thorax) within each frame using a modified version of LEAP (Pereira et al., 2019) (544 instances were used for training; Fig. S1Ai-ii and Supp Movie S1). Using the same software and neural network architecture, a separate network was then trained to group these detections together with the correct animals by inferring part affinity fields (Fig. S1Aiii; Cao et al., 2017). This enabled estimation of the vector that represents fly heading for both flies (Fig. S1Aiv).

### Linear classification of single frames

17 parameters were extracted for each frame based on the tracking of male and female centroid and heading (Fig. 2A), describing either the female movements (fFV – female forward velocity; fLS – female lateral speed; fFA/fLA – female forward/lateral acceleration; fRS – female rotational speed), male movements (similar female movements – mFV/mLS/mFA/mLA/mRS), or male-female interaction (mfDist – male-female distance; fmAngle – female heading relative to the female-male axis, mfAngle – male heading relative to male-female axis; fmFV or fmLS – female speed in the male direction or in the perpendicular axis; mfFV or mfLS – male speed in the female direction or perpendicular). Using 17 parameters for each frame, we trained binary support vector machine (SVM) linear classifiers to find the parameters (dimensions) that best separate between the groups. We first trained classifiers that separate between frames that belong to experimental flies (class 1, pC1-Int activated, either one condition - d0/d3/d6 or all groups together, d0-d6), and controls (class 2). We trained 90 classifiers, randomly choosing a set of 3000 frames from each class (‘training set’; non-overlapping - the same frame was never used in two classifiers; increasing the number of frames beyond 3000 did not increase performance). We used the MATLAB R2019b procedure *fitcsvm* (MathWorks, Natick, MA), with a linear kernel. We then used a separate set of 30,000 frames per class for each classifier (‘validation set; the same frame was never used twice, either between classifiers or between sets) to test the performance of each classifier (fraction of frames correctly classified). We then choose the 30 best-performing classifiers (Fig. 2B for control vs d0-d6). We used a third set of frames for each classifier (30,000 frames/class, again – with no overlap with other sets) to measure the performance of each classifier. The MATLAB function *predict* was used to find the SVM-predicted class for each frame in the validation or train set. Performance was calculated as the percent of frames correctly classified (Fig. 2C). For each weight (out of the 17; control vs d0-d6), we looked at the distribution coming from the 30 independent classifiers, and tested whether the mean was significantly different than zero (Fig. S1E).

We used a two-sample t-test to measure the probability that the mean weight associated with each parameter is different from zero (Fig. S1E). We found 8 out of the 17 parameters to be highly significant (*P<0.0001 in S1E).

### Clustering behaviors based on single frames

The 8 most significant parameters found by the SVM classifier (see previous section) were used for classification. We took the same number of frames from each group (control/d0/d3/d6) - 357997 frames (99.4 minutes) per group, corresponding to the number of frames in the smallest group (d6). We repeated the clustering 30 times (Fig. 2D, black dots), each time selecting 1hr39min of data from each one of the other groups (d0, d3, d6, control) randomly (with replacements – the same frame could be used in multiple repeats), therefore having >1.4 million frames for clustering on each repeat. The sets are not independent (overlapping frames between repeats) and no statistical test was performed over the repeats. After z-scoring each parameter (over all the frames in a given repeat), k-means clustering was performed (using MATLAB function *kmeans*), allowing 20 clusters and a maximum of 500 iterations (other parameters set to default). We found that the first 7 largest clusters (cluster size being the number of frames in the cluster) capture 90.4% of the frames, averaged over repeats (Fig. 2D, inset). To match clusters between repeats (for each cluster number in repeat 1, find the corresponding cluster number in repeats 2-30), we used the smallest distance between clusters, by calculating the mean square error over the weights (the variability in weight size across repeats is shown in Fig. S1G).

### Machine learning based classification of behavioral epochs

The Janelia Automatic Animal Behavior Annotator (JAABA; http://jaaba.sourceforge.net/) was used to detect epochs of ‘female shoving’ and ‘female chasing’. Two independent classifiers (‘shoving classifier’, ‘chasing classifier’) were trained, one for each behavior. We used the automatic segmentations to find examples for shoving and chasing epochs, used as a first step in training each classifier. We then added example epochs (positive and negative examples are used for each classifier), in an iterative manner (using examples where the classifiers made wrong predictions). Altogether we used 24,222 frames (6.7 minutes) to train the ‘shoving classifier’, and 11,941 frames (3.3 minutes) for the ‘chasing classifier’.

The classifiers was based on the 17 parameters defined above (denoted as ‘per-frame’ features), as well as on ‘window features’ (‘mean’, ‘min’, ‘max’, ‘change’, ‘std’, ‘diff_neighbor_mean’, ‘diff_neighbor_min’, ‘diff_neighbor_max’, ‘zscore_neighbors’ with a window radius of 10 and default ‘windows parameters’), therefore taking into account longer timescales for classification, rather than the single frames we used for SVM classification and k-means clustering (see Fig. S1H for comparison). We cross validated each classifier before applying the classification on all the data, using the cross-validation procedure available in JAABA package (with default parameters). 94.2% of the frames annotated by the user as shoving were correctly classified as shoving, while 92.8% of the frames annotated as no shoving were classified as no-shoving. For the ‘chasing classifier’, we got 96% and 90.8% success in classifying chasing and no-chasing. The trained shoving classifier was used to annotate each frame as belonging or not belonging to ‘female shoving’ epoch, and the trained chasing classifier was used independently to classify each frame as belonging or not-belonging to a ‘female chasing’ epoch.

### Manual tracking

We tracked a subset of the data manually (pC1-Int, d0-d6), to confirm our automatic behavior detection, as well as in search for more rare events, or events that are not captured due to tracking issues. Three behaviors were annotated by two observers: female shoving, female chasing and circling (Fig. S2B,D). The two observers annotated different sets of movies, while a small subset (N = 5 movies) were annotated by the two observers and we confirmed that both detected three behaviors (shoving, chasing, circling) similarly. Female circling was not detected by our automated procedures for two reasons. First, during circling male and female bodies often overlap, causing large errors in heading detection. Second, these events are relatively sparse. One observer also detected three other rare behaviors: head butting, female mounting (Fig. S2E, Supp Movie S3) and wing extension (Fig. S2F-G, Supp Movie S2).

### Statistical analysis

Statistical analysis was performed using MATLAB (Mathworks, Natick, MA) procedures, and corrected for multiple comparisons using the Bonferroni correction when appropriate. The details on the statistical test used are listed under the Results section and the Figure legends. Black lines between two groups indicate a statistically significant difference between the groups after multiple comparison correction, while a red line indicates that the difference is statistically significant only when multiple comparisons test is not used. To test for significant differences in copulation rate, we used Cox’s proportional hazards regression model, using the MATLAB procedure *coxphfit*. ‘Censoring’ was used to account for the fact that some flies copulated within the 30-minute time window (after which the experiment was terminated), while others did not. The correlation between female velocity and male song (Fig. 1D,G, S1D) was done as previously described (Clemens et al., 2015). Briefly, female absolute speed and male song were averaged over 1-minute windows. In each window we calculated the mean value of female (absolute) speed, bout amount (the total amount of song in the window), bout number (the number of song bouts in the window) and bout duration (the mean bout duration in the window). Then, for each condition, we calculated the correlation between female speed and male song by pooling all windows for a given group together. The MATLAB procedure *corr* was used to calculate the Pearson correlation, and one way analysis of covariance (ANOCOVA) was used to compare the slopes (x,y being the male song and the female speed) between groups using aoctool (MATLAB). The 30 SVM (Support Vector Machine) classifiers (Fig. 2B-C, S1E) were trained using non-overlapping sets of frames and are therefore considered independent. One-sample t-test was used to calculate a test decision for the null hypothesis that the 30 weight values (for a given parameter) come from a normal distribution with a mean of zero (and unknown variance). For each parameter, −log(P) is shown, and a vertical dashed indicates P<10^-4^ (Fig. S1E).

### *In vivo* whole-brain calcium imaging

We imaged brain activity following pC1 optogenetic activation (though the microscope objective) under a two-photon custom made microscope (Pacheco et al., 2019) in females, using the calcium indicator GCaMP6s (Chen et al., 2013). Both GCaMP6s and the structural marker tdTomato (Shaner et al., 2004) were expressed pan-neuronally in blind flies (NorpA[36] mutant) using the nsyb enhancer (Bussell et al., 2014). For pC1 activation, we used the same temporal pattern as the one used in the behavioral experiments: 5 minutes of light on, at 100Hz and 50% duty cycle. Imaging started 5 minutes before stimulus onset, where baseline activity was measured, and lasted 9.5 minutes after stimulus offset for whole-brain imaging and 30 minutes after stimulus offset for *doublesex* imaging. While the red shifted channelrhodopsin ReachR was used for behavior experiments to minimize optogenetic activation by background light, we used Chrimson for two-photon imaging for two reasons. First, to minimize the amount of bleed-through from the optogenetic activation light to the green photomultiplier tube (PMT). For this reason we also choose a longer wavelength of 700nm (M700L4, Thorlabs, Newton, NJ with a band pass filter FF01-708/75-25, Semrock, Rochester, NY) that is well separated from the range of light that cross the green PMT entrance filter (Semrock FF01-593/40-25). Second, we wanted to minimize the amount of light needed for neuronal activation by using a more sensitive effector, to reduce the amount of heat accumulating in the brain during imaging (on top of the heating caused by the two photon laser, whose power was limited to 15mW). We are aware of possible differences in pC1 activation level between the behavioral and imaging experiments. Based on existing literature, we tried to choose an activation level for the imaging experiments that will roughly match the activation induced during behavior. In order to match the activation level between the behavioral experiments (ReachR, 627nm light, intact fly) and the imaging experiments (csChrimson, 700nm light, cuticle removed above the fly brain) we used data from existing literature. By comparing the amount of light needed for driving proboscis extension reflex (PER) in 100% of adult flies in (Inagaki et al., 2014) (ReachR, 1.1mW/mm^2^, 627nm) to the level of light used to saturate PER score in (Klapoetke et al., 2014) (CsChrimson, 0.07mW/mm^2^, 720nm), taking into account the different duty cycles used in the two studies and given the penetration rate through the cuticle (Based on (Inagaki et al., 2014) Fig. 1A, around 6% of the light penetrates at 627nm), we choose a light intensity of 0.013 mw/mm^2^.

A volume of ∼307×307×200 µm^3^ from the dorsal part of the central brain was scanned at 0.1 Hz (1.4×1.2×2 µm^3^ voxel size), covering a complete dorsal quadrant (full anterior-posterior axis of the central brain) which represents about 58.02 +/- 3.97 % of the whole hemisphere (mean +/- SD, N = 28 animals). Volumetric data was processed as described in (Pacheco et al., 2019). In brief, tdTomato signal was used to motion-correct volumetric time-series of GCaMP6s signal in XYZ axis (using the NoRMCorre algorithm (Pnevmatikakis and Giovannucci, 2017)). Volumes were spatially resampled to have isotropic XY voxel size of 1.2×1.2×2 µm^3^ (bilinear interpolation on X and Y axes), and temporally resampled to correct for different slice timing across planes of the same volume, and to align timestamps of volumes relative to the start of the optogenetic stimulation (linear interpolation). Next, the GCaMP6s signal was 3D-ROI segmented to obtain spatial and temporal components per segmented ROI using CaImAn (Giovannucci et al., 2019; Pacheco et al., 2019). Code to perform these processing steps are available at https://github.com/dpacheco0921/CaImProPi. In addition, to further remove residual motion artifacts from the GCaMP6s signal, in particular slow drift over tens of minutes, we performed independent component analysis (ICA) on the tdtomato (F_tdtomato_) and GCaMP6s (F_GCAMP_) signal for each ROI independently, similar to (Scholz et al., 2018). To remove opto-related artifact bleeding through the red channel, F_tdtomato_ was linearly interpolated from 20 seconds before stimulus onset to 20 after stimulus offset (to ignore opto-related artifact bleeding through the red channel) and random noise (from normal distribution centered at 0) added to interpolated timepoints. F_tdtomato_ was then smoothed (moving average with a window of 50s), and ICA was used (rica function implemented in MATLAB) to extract background and signal components from F_tdtomato_ and F_GCAMP_. Independent component highly correlated to F_tdtomato_ (absolute correlation coefficient > 0.9) was considered the background component (ICA_Background_(F_tdtomato,_ F_GCAMP_)), while the other component considered the signal component (ICA_Signal_(F_tdtomato,_ F_GCAMP_)). Sign of ICA_Signal_(F_tdtomato,_ F_GCAMP_) was corrected using the sign of the correlation between ICA_Signal_(F_tdtomato,_ F_GCAMP_) and F_GCAMP_. For ROIs extracted from pan-neuronal data we report calcium signals as ICA_Signal_(F_tdtomato,_ F_GCAMP_) as shown in Figure 5B.

We defined responsive ROIs as ROIs with a mean activity during optogenetic stimulation (Ft1) higher than 3***σ***_o_ (***σ***_o_ - standard deviation of activity during baseline). We then split ROIs into transiently and persistently active units using the mean activity after optogenetic stimulation (Ft2, from stimulus offset to 5 minutes after stimulus offset), transient ROIs had Ft2 ≤ 3***σ***_o_, while persistent ROIs had Ft2 > 3***σ***_o_. To evaluate the diversity of these coarse activity types, we hierarchically clustered transient and persistent responses (we evaluated the number of clusters these response types split into using the consensus across Calinski-Harabasz, Silhouette, Gap, and Davies-Bouldin criteria), obtaining 2 clusters of transient responses and 2 clusters of persistent responses (Figs. 5C-D).

For recordings of Dsx+ cell types, we imaged pC1, pC2, pCd1, pCd2, and pMN2 cells (1-2 groups at a time), located in the dorsal side of the central brain, at a speed of 0.5-0.25 Hz (0.5×0.5×1 µm^3^ - 2.5×2.5×1 µm^3^ voxel size). Volumetric time-series of GCaMP6s signal was motion-corrected in the XYZ axes (using the NoRMCorre algorithm (Pnevmatikakis and Giovannucci, 2017)), and temporally resampled to correct for different slice timing across planes of the same volume, and to align timestamps of volumes relative to the start of the optogenetic stimulation (linear interpolation). Dsx+ somas were manually segmented by finding the center and edge of each cell body stack by stack (Deutsch et al., 2019)).

### Immunostaining

Flies were dissected in S2 insect medium (Sigma #S0146). Dissected brains were moved through 6 wells (12ηl/well) containing a fixation solution (4% paraformaldehyde, Electron microscopy sciences #15713 in PBT (0.3% Triton in PBSX1; Triton X-100 Sigma Aldrich #X100; PBS - Cellgro #21-040), before sitting for 30 minutes on a rotator at room temperature. Following fixation, brains were moved through 6 wells containing PBT, 15 minutes in each well. Then, brains were transferred through 4 wells containing a blocking solution (5% Goat Serum in PBT; Life Technologies #16210-064), and sitting in the last well for 30 minutes. Brains were then moved to a solution containing primary antibodies (see below) and then incubated for two nights at 4C (sealed and light protected). After 8 washes (20 minutes per wash) in PBT, brains were incubated overnight with secondary antibodies. After 8 washed (20 minutes each) in PBT, brains were placed on a slide (Fisher Scientific #12-550-15), between two zero numbered coverslips used as spacers (Fisher Scientific 12-540B) and under a coverslip (Fisher Scientific #12-542B), and Vectashield (Vector Laboratories) was applied. Nail polish was used to seal around the center coverslip edges, and brains were stored in dark at 4C overnight to harden, before imaging. The primary antibody solution contained 2 primary antibodies in blocking solution: mouse nc82 (anti-Bruchpilot; mAb DSHS - developmental studies Hybridoma bank; 1:20), chicken anti-GFP (Invitrogen A11039; 1:2000). Secondary antibody solution contained 2 secondary antibodies in blocking solution: Goat anti-mouse Alexa-Flour (AF) 568 (Invitrogen A-11004, 1:400), Goat anti-chicken AF-488 (Invitrogen A11039; 1:300). Imaging was done using a Leica confocal microscope (TCS SP8 X). Fig. 1A was modified from (Deutsch et al., 2019).

### Identification and proofreading of neurons in FlyWire

Neurons in a complete EM volume of an adult female brain (Zheng et al., 2018) were automatically reconstructed in FlyWire (flywire.ai, manuscript in preparation). Within FlyWire, we first searched for reconstructed segments that match the morphology of known pC1 cells. We used anatomical landmarks to find the bundle that projects dorsally from pC1 cells bodies (Fig. 1A, red arrow). We then looked at 2 cross sections of this bundle in each hemisphere (Figs. 3B and S3A) and looked systematically at all the segments that pass through this bundle. Based on known morphology of female pC1 cells (Deutsch et al., 2019; Kimura et al., 2015; Zhou et al., 2014), we defined cells as pC1 when they crossed through the pC1 bundle, and also projected to the lateral junction (Fig. 1A). Similarly, pC2l cells were found by looking through the pC2l bundle (Fig. 1A, green arrow; we refer to these cells as pC2 in this paper). The aIP-g cells were first found by searching for neurons synaptically connected to pC1-Alpha-l (see below), and other aIP-g cells were found by systematically exploring a cross section within the aIP-g bundle (Figs. 6E and S5B). pMN1 and pMN2 were found when mapping the pC1-Alpha-l synaptic partners, and then named pMN1 and pMN2 based on their morphology (Deutsch et al., 2019; Kimura et al., 2015). pC1 and pC2 cells were sorted manually into subtypes based on morphology (Fig. 3C and S5E).

Proofreading of a neuron was performed using the tools available in FlyWire (flywire.ai, manuscript in preparation). In short, this process has two parts: (1) removing (‘splitting’) parts that do not belong to the cell (‘mergers’), such as parts of glia or parts of other neurons (for example, when detecting two cell bodies in one segment), and (2) adding missing parts (‘merging’). We had an average of 5.4 splits and 10.7 merges per neuron, and proofreading a single cell took 43 minutes on average (we measured the proofreading time for a subset of the cells we proofread). Proofreading was complete when no additional obvious mergers were found, and we couldn’t identify missing parts at the edge of any processes. In some cases, the known morphology of the cell (e.g., pMN1 or pMN2) or the existence of other cells with similar morphology (in the same or the other hemisphere) were used to verify that no major processes were missing. Sorting cells into types was done manually, based on their morphology. pC1 was divided into seven subtypes, pC2 and aIP-g were divided into three subtypes each.

Assigning names to known neurons we found in the EM volume was done solely based on morphology. It is possible, that in some cases (e.g., for pC2 or pC1 cells), some of the neurons we found are not actually Dsx+ cells. More work is needed to compare LM based and EM based morphologies, and to classify cell types based both on morphology and connectivity (Xu et al., 2020).

### Mapping synaptic inputs and outputs in FlyWire

We mapped all the direct inputs and outputs of a single pC1-Alpha-l neuron (Fig. 3C) by manually detecting pre- and postsynaptic partners for this cell. After proofreading the cell (see details above), we looked systematically, branch by branch, for synaptic partners based on previously defined criteria. For a contact to be defined as a chemical synapse, it had to meet three conditions: (1) the presence of a synaptic cleft between the pre- and postsynaptic cells, (2) presynaptic active zone with vesicles near the contact point, and (3) one of two (or both) must exist: a presynaptic T-bar adjacent to the cleft (Fouquet et al., 2009) at the presynaptic terminal or a postsynaptic density (PSD, (Ziff, 1997)). In flies, PSDs are variable, and are often unclear or absent (Prokop and Meinertzhagen, 2006). Typically, we observed T-bars rather than PSDs, as a T-bar is easier to identify. Our criteria was slightly more conservative than the one used in (Zheng et al., 2018), possibly leading to less false positives (wrongly assigned synapses), and more false negatives (missing synapses). Once a synapse was detected, we then looked for the post-synaptic partner. Around 10% of the inputs to pC1-Alpha-l and about 60% of the outputs were short segments (‘twigs’), that we could not connect to backbones in order to identify or proofread the connected neuron. The twigs were not restricted to a specific part of the pC1-Alpha-l cell, and we therefore believe that they do not impose a bias on the distribution of pC1-Alpha-l connections, though it is possible that specific output types (e.g., cells with thinner processes) are less likely to be detected.

Following the detection of pC1-Alpha-l synaptic partners, we mapped the inputs and outputs to pC1-Alpha-l in three steps. First, we manually proofread the input and output segments. Second, we eliminated cells that connect to pC1-Alpha-l with less than 3 synapses, to reduce the number of potential false positives, and to focus on stronger connections. We ended up having 78 input and 52 output cells. Third, we sorted cells manually into cell types based on morphology. Some cells were classified based on known morphology from light microscopy (pMN1, pMN2, pC1, pC2, aIP-g). In order to look for connections between pC1 and pC2 cells (the largest sets of Dsx+ neurons) in an unbiased way (not focusing on specific types or individual pC1 or pC2), we first identified and proofread pC1 and pC2 cells. Synaptic connections between individual cells of pC1 or pC2 type were detected by manually inspecting the volume plane by plane. Once a pair of segments that came within proximity of one another was detected, we zoomed and looked for synaptic connections based on the criteria defined above.

Finding the best match in the single clone dataset FlyCircuit for a given EM segment was done in two steps. First, an .swc file was generated for a given segment (using the automatically segmented cells rather than the proofread ones for technical reasons). Second, we performed an NBLAST search (Costa et al., 2016) either online (http://nblast.virtualflybrain.org:8080/NBLAST_on-the-fly/) or using ‘natverse’, an R package for neuroanatomical data analysis (Manton et al., 2019). For visualization purposes, we first created mesh files (.obj) for proofread neurons, and then used either Meshlab (http://www.meshlab.net/) to create images, or Blender (https://www.blender.org/) to create movies (see support/FAQ in https://flywire.ai/ for instructions on creating .swc and .obj files).

---

**Supplemental Table 1:**
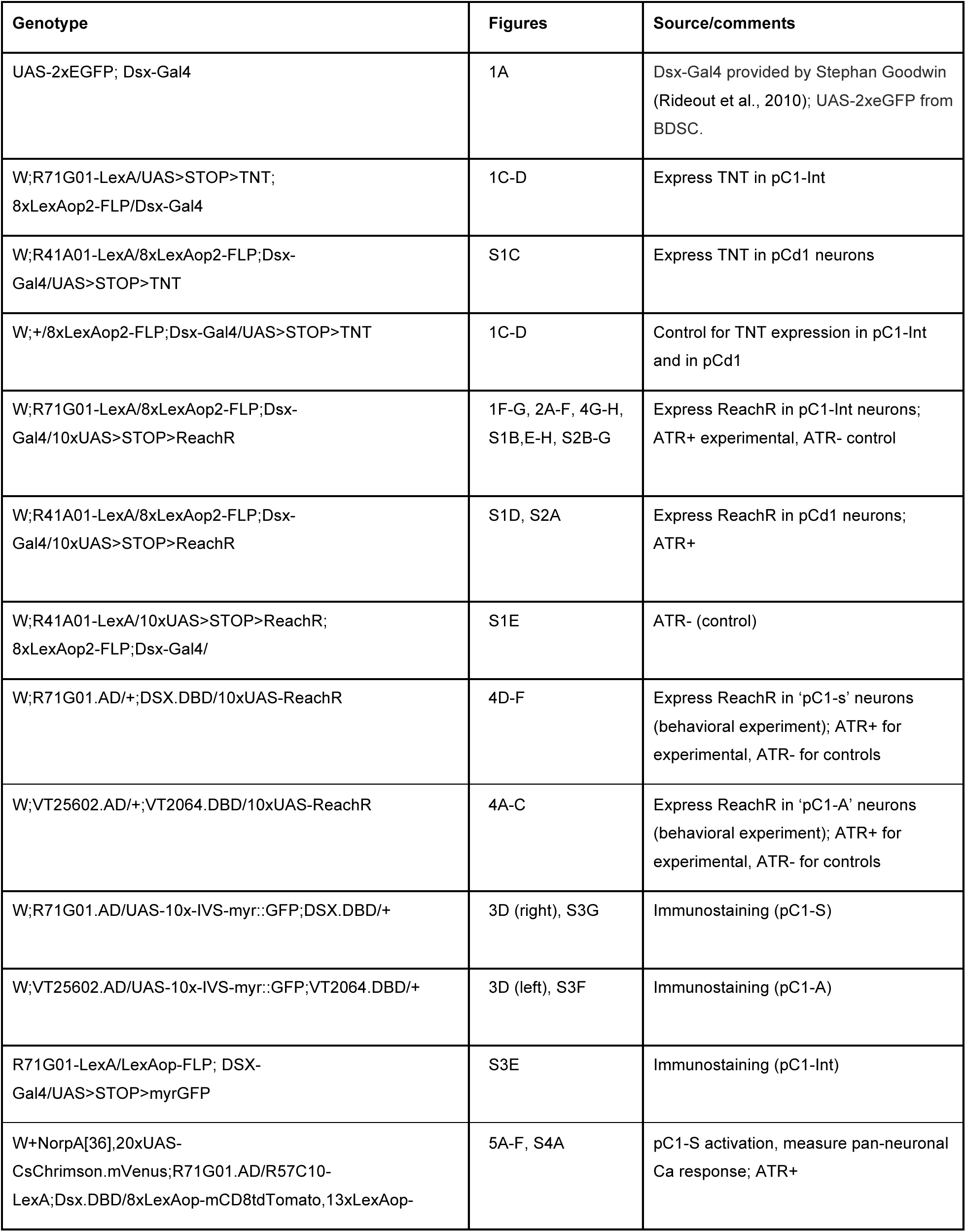
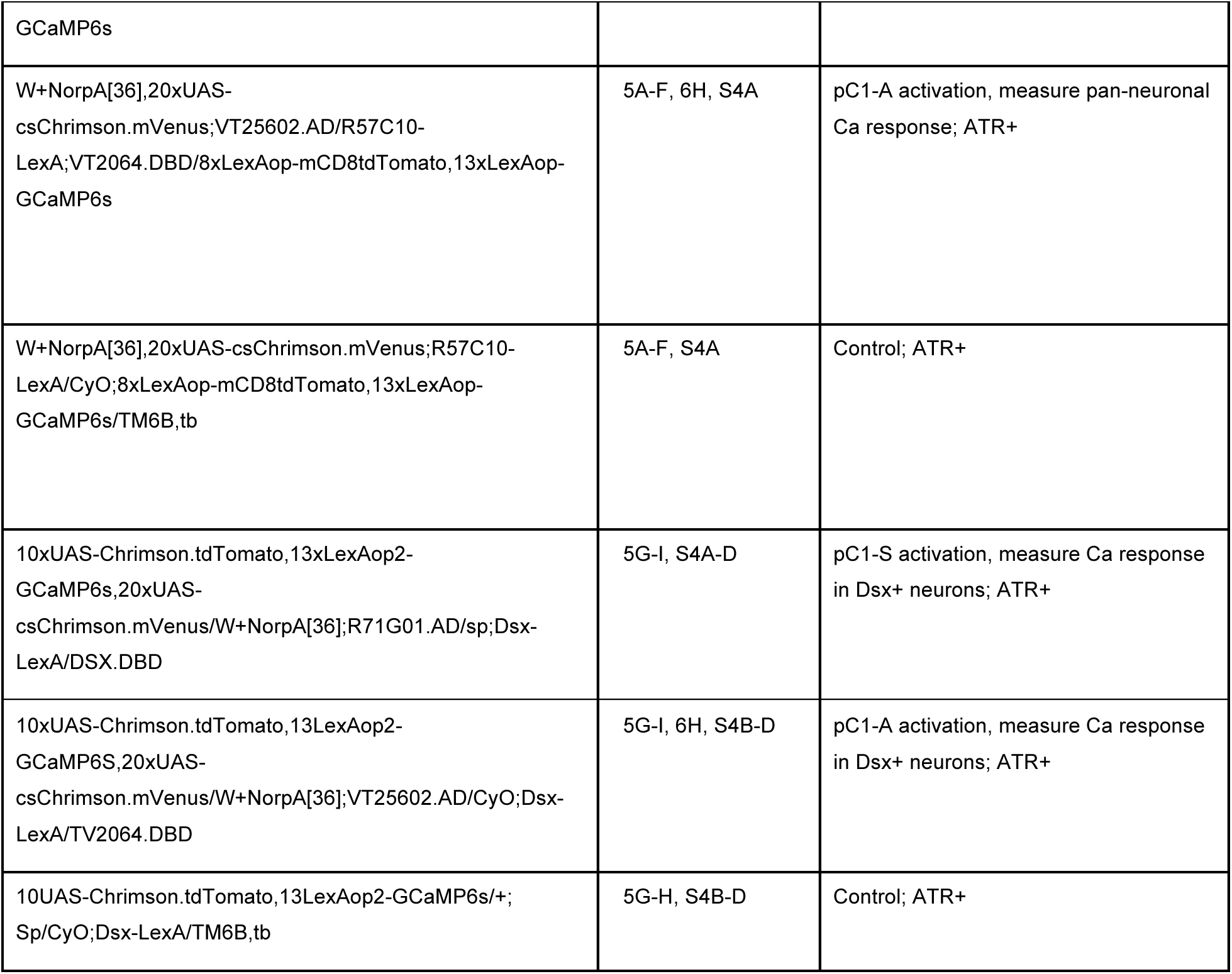
Genotypes used in this study

## Supplementary Figures

**Figure S1.**
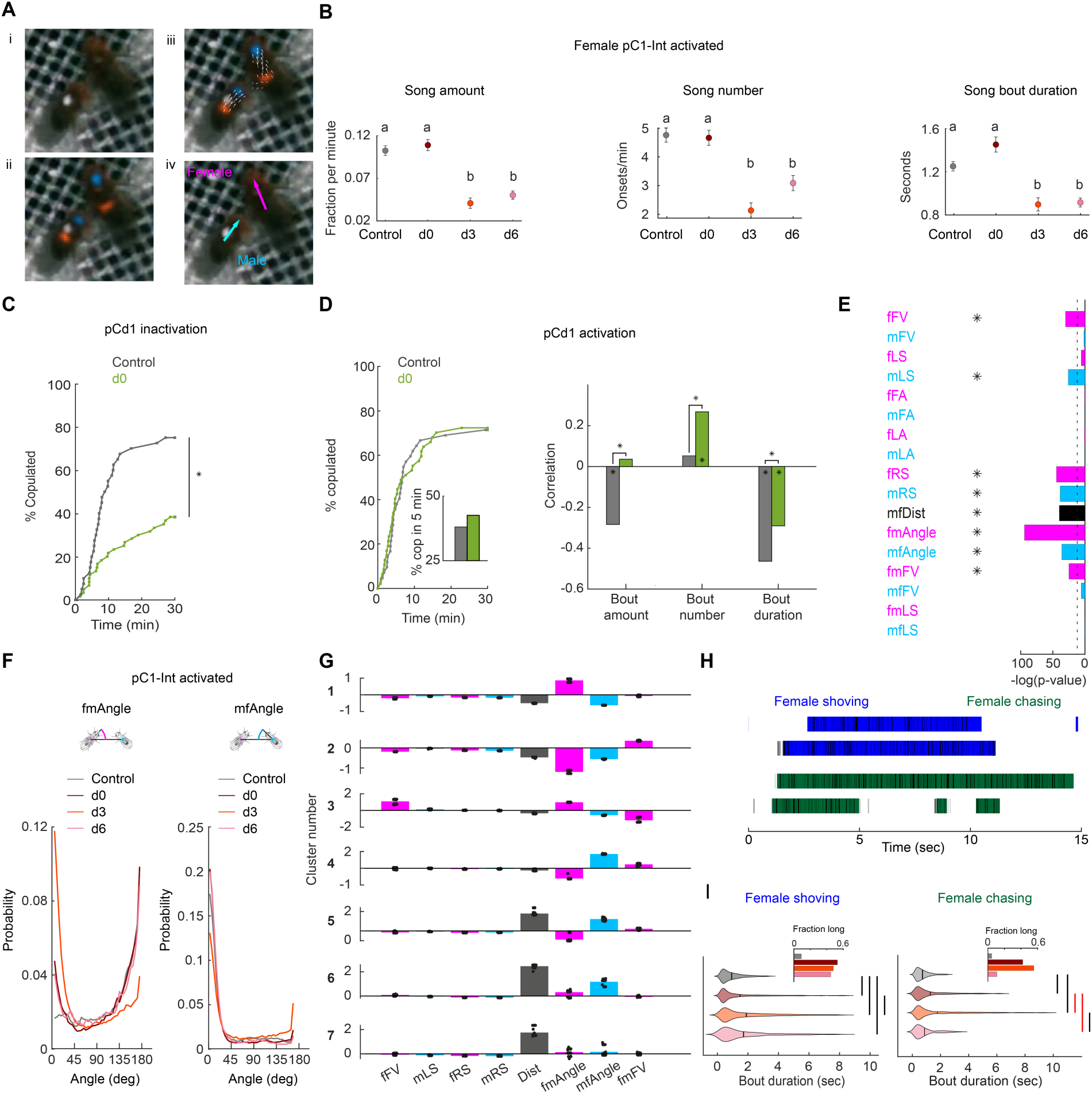
(related to Figures 1 and 2) **(A)** (i) A single video frame of a male (with painted dot) and a female in the behavioral chamber. (ii) Confidence maps (Pereira et al., 2019) for male and female head (blue) and thorax (red). (iii) Part affinity vector fields (Cao et al., 2017). (iV) Heading of male (cyan) and female (magenta). **(B)** Song amount (left), number (middle) and duration are defined as in as in (Clemens et al., 2015), see Methods. Mean and standard error over 1-minute windows are shown for each condition. Mean and standard errors are shown. Statistically different groups (two sample t-test) are marked with different letters (a, b). P<0.0003 for all pairs of groups marked (a, b) for a given song parameter. **(C)** Percent of pairs that copulated as a function of time (P = 0.0001, Cox proportional hazards regression, see Methods; N = 40, 60 pairs for Control, d0). pC1d>TNT (see Table 1 for full genotype). **(D) Left:** Same as (C) for pCd1 activated female (pCd1>ReaChR, see Table 1 for full genotype). P = 0.93, Cox proportional hazards regression, see Methods; N = 42, 47 pairs for Control, d0. **Inset:** percent of flies copulated in 5 minutes. **Right:** Correlation between song parameters and female speed, averaged over 1 minute windows as in (Clemens et al., 2015) (see Methods). Significance (indicated by an asterisk above the line connecting a pair of groups), was measured using ANOCOVA (MATLAB *aoctool*) and multiple comparison correction (*P<0.01). An asterisk in the base on a bar indicates a significant correlation (MATLAB corr, *P<0.01). **(E)** Bar height indicates −log(P-value) for the probability that the mean distribution of SVM (Support Vector Machine) weights (over 30 independent classifiers) associated with each weight (Figure 2B) is significantly different than zero. Natural log is used. Dashed line indicates P-value = 10^-4^. Asterisks indicate weights associated with distributions with P-value < 10^-4^. **(F)** Distribution of fmAngle and mfAngle (Fig. 2A) are shown for 4 experimental conditions (4.5 deg bin size). fmAngle/mfAngle are the absolute number of degrees the female/male needs to turn in order to point to the centroid of the other fly (see cartoons). **(G)** The weights associated with each behavioral cluster (Fig. 2D), for the 8 significant weights (Supp Fig. S1E) that were used for clustering. Each dot represents a single clustering repeat (see Methods). **(H)** Frames that belong to the shoving (blue) or chasing (green) behavioral clusters (Figure 2D) are indicated as black horizontal lines. JAABA classification for the same 15 seconds is indicated as horizontal bars. **(I)** Violin plots (MATLAB violin) are shown for bout duration of female shoving (left) and chasing (right) bouts based on JAABA classification. Means are shown as black lines (0.99/1.47/1.88/1.7 seconds for control/d0/d3/d6). Black vertical line indicates a significant difference between groups (p<0.05, two sample t-test). Red line indicates that the difference is significant only if multiple comparison correction is not applied. **Inset:** The fraction of all frames in the experiment that belong to long bouts (≥5 seconds). In the main plots (but not in the insets and not for statistical measures) the smallest and largest 5% bout durations were excluded for each condition.

**Figure S2.**
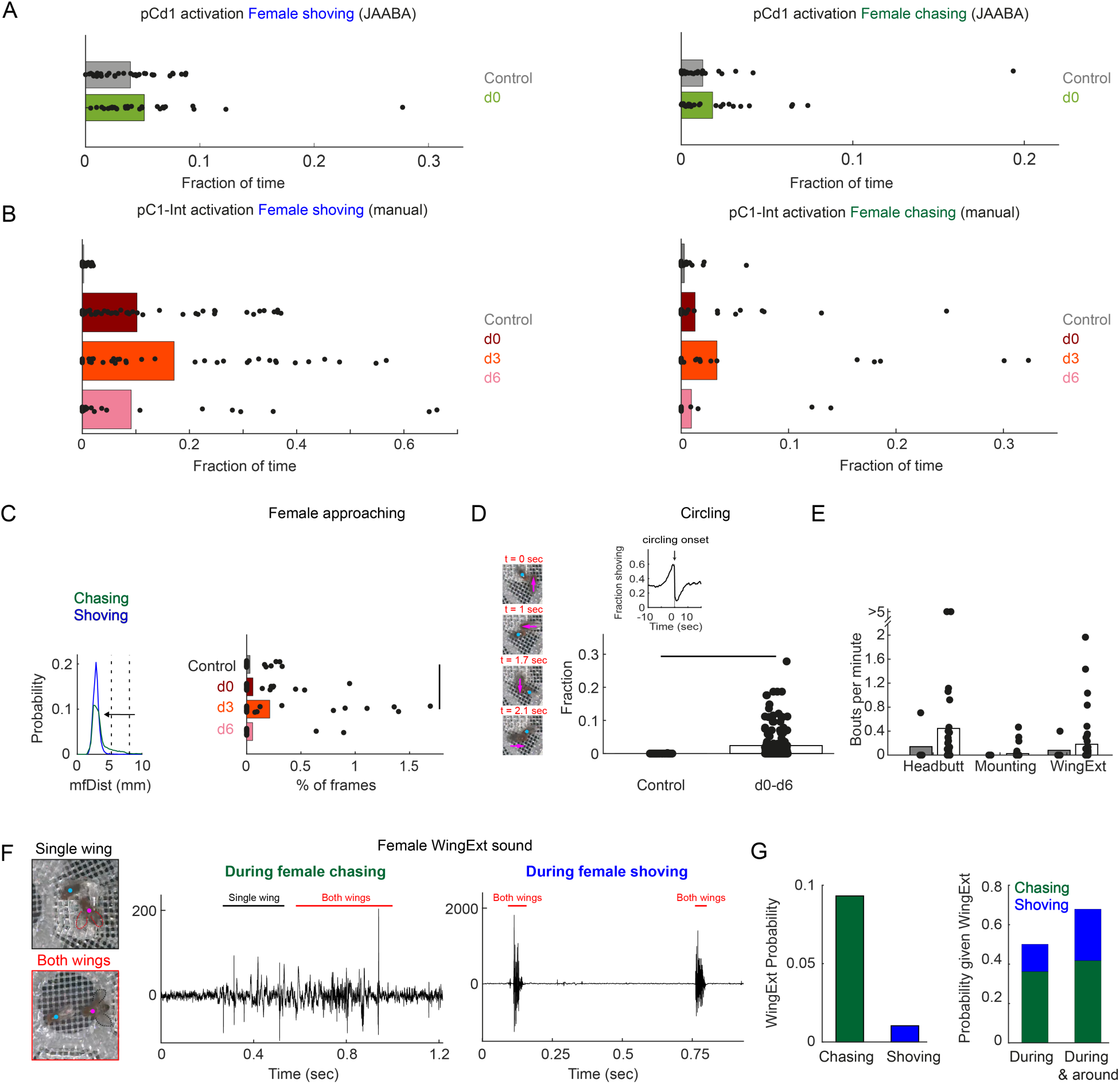
(related to Figure 2) **(A)** Fraction of time the female spent shoving (left) or chasing (right) following pCd1 activation, based on JAABA classification. **(B)** Fraction of time the female spent shoving or chasing following pC1-Int activation, based on manual scoring. **(C) Left:** Distribution of mfDist (male-female distance) during female chasing (green) and shoving (blue) for d0-d6 conditions. The horizontal arrow illustrates the criterion used for defining ‘female approaching’ events: the female is approaching the male from large distance (> 98 percentile mfDist during shoving or chasing) to short distance (< 95 percentile for distance mfDist during shoving/chasing), while continuously heading towards the male (fmAngle < 30 deg). **Right:** The percent of frames for each condition that belong to ‘Female approaching’ epochs. Black line indicates significant difference (P < 0.05, two sample t-test with Bonferroni correction for multiple comparisons). **(D) Left:** Four example frames from a single ‘circling’ epoch, separated by 90 deg in the female heading direction (see also Supp Movie S3). In this example, a female completed 270 deg in 2.1 seconds. **Right:** Fraction of time the male and female are spending ‘circling’ based on manual annotation. The difference was statistically significant between the control and conditions d0-d6 taken together (p<0.05, two sample t-test), but not when taking each condition alone. **Inset:** The fraction of time the female spent shoving (d0-d6) aligned to circling onsets indicates high probability for shoving shortly before circling onset. **(E)** Number of bouts per minute are shown for manually detected behaviors: ‘female headbutting’, ‘female mounting’ and female extending one or two wings (see Supp Movies 2,3). The two points with >5 represent 5.5 and 8.1 bouts per minute. **(F)** Example frames with female unilateral (top) or bilateral (bottom) wing extension (WingExt). **Middle**: example sound trace during female chasing with unilateral and bilateral wing extensions. **Right**: example sound trace during female shoving with bilateral wing extension. Note that the sound evoked by female wing extension during shoving was an order of magnitude larger than the sound evoked when the female extended one or two wings during chasing. **(G) Left:** Wing extension was manually detected in 9.3% of all frames (d0-d6 taken together) during chasing epochs, and in 1% of the frames during shoving epochs. **Right:** 50% of the frames detected as ‘wing extension’ were part of female chasing or shoving, and 67.9% of the frames with wing extension occurred during or around chasing or shoving bouts (‘around’: 2 seconds before epoch onset until 2 seconds after epoch offset).

**Figure S3.**
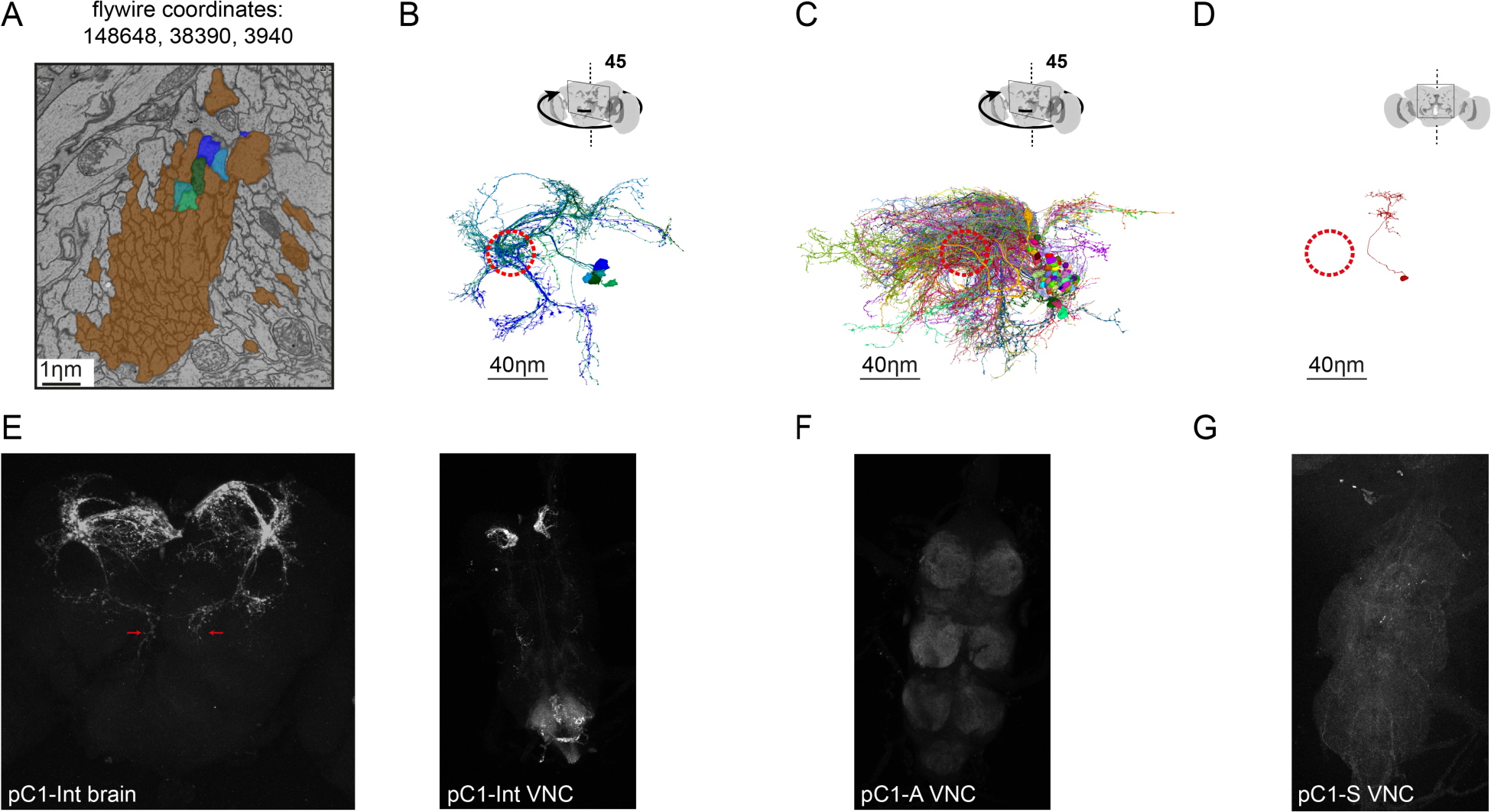
(related to Figures 3,4) **(A)** Cross section of left pC1-bundle. All segments going through this cross-section or through another cross section (separated by 140 slices, equal to 5.6 µm; FlyWire coordinates: 148388, 39874, 4080) were tested. Similarly, two cross sections were tested in the right hemisphere. Automatically reconstructed segments crossing any of these 4 cross sections were sorted based on morphology. Segments that included a projection to the lateral junction (colored in blue/green/cyan) or segments that were too short to judge were proofread, and considered pC1-like neurons. Neurons that passed through any of the cross sections, but did not project to the lateral junction (colored in brown) were not further analyzed. **(B)** Proofread neurons that go through the cross section in (A) and also project to the lateral junction. Dashed red circle marks the lateral junction (Cachero et al., 2010). **(C)** Neurons that go through the cross section in (A), and do not project to the lateral junction. Dashed red circle marks the lateral junction location. **(D)** The most common type of neuron from those shown in (C). **(E)** pC1-Int neurons expressing GFP (see Table 1 for full genotype). **Left:** A maximum z-projection is shown for the pC1-Int processes. Red arrows mark the medial projections (left and right hemispheres) that exist in pC1-Alpha, but not in other pC1 cells types. **Right:** pC1-Int expression in the female VNC. **(F)** pC1-A in the female VNC expressing GFP (see Table 1 for full genotype). **(G)** pC1-S in the female VNC expressing GFP (see Table 1 for full genotype).

**Figure S4.**
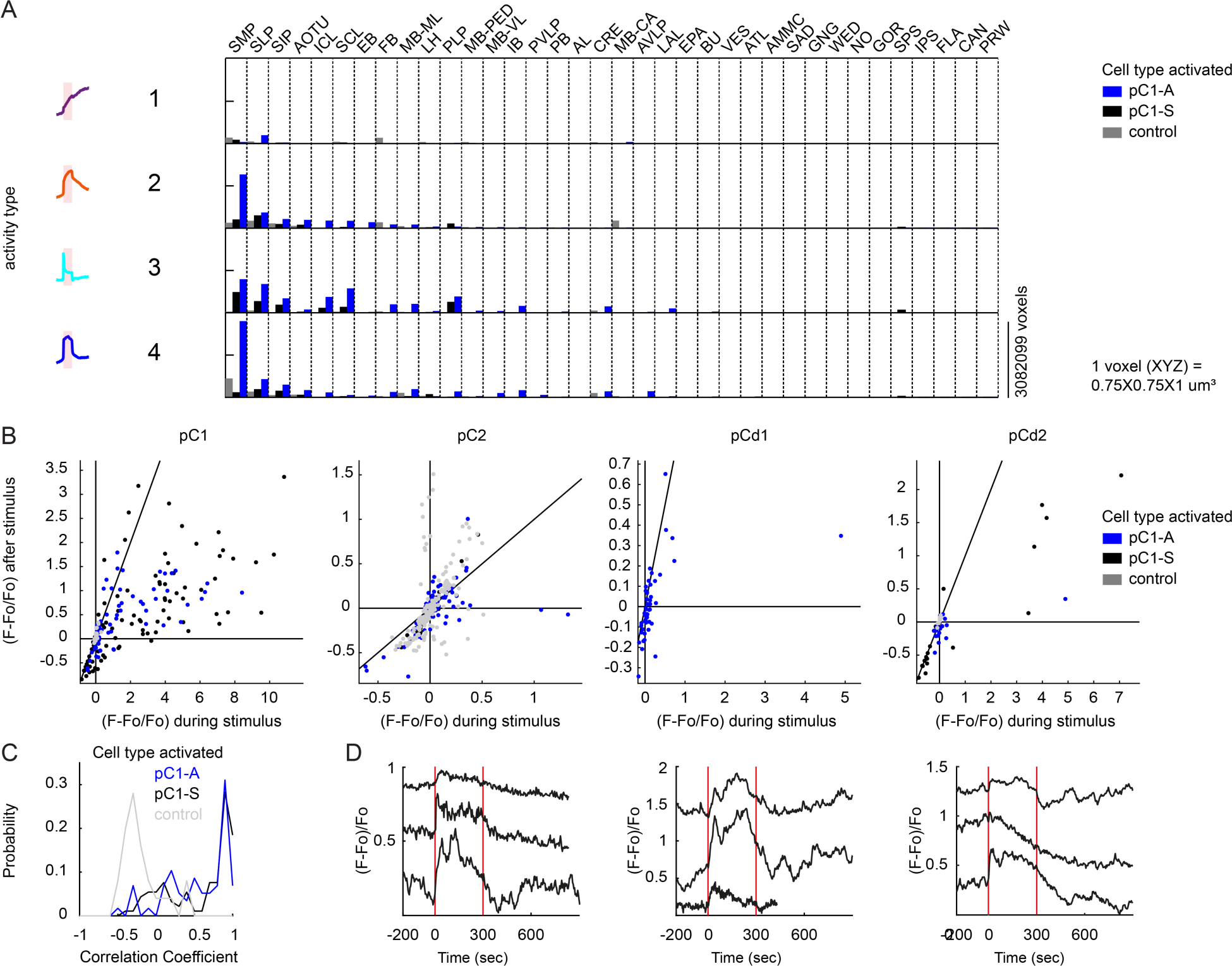
(related to Figure 5) **(A)** Average number of voxels occupied by all ROIs belonging to each response type (as in Fig. 5F) across all 36 central brain neuropils for each condition (pC1-A or pC1-S activation, or controls), sorted from left to right by the amount of response type 2. **(B)** Mean calcium responses during t1 (x-axis) versus during t2 (y-axis) as in Figure 5H for the major Dsx+ cell types (pC1, pC2, pCd1, and pCd2). Activity units are in (F - Fo)/Fo, where Fo is the mean activity during baseline, and F is the mean activity during t1 or t2 for x and y axes, respectively. Each dot represents a single cell, and dot colors refer to different conditions (pC1-A or pC1-S activation, or controls). **(C)** Distribution of correlation coefficient of pC1 cell responses to optogenetic stimulation for all conditions (pC1-A or pC1-S activation, or controls). **(D)** Example traces of pC2, pCd1 and pCd2 cells with stimulus-locked transient responses (traces were selected based on the correlation of the cell response to stimulus presentation, correlation coefficient > 0.5).

**Figure S5.**
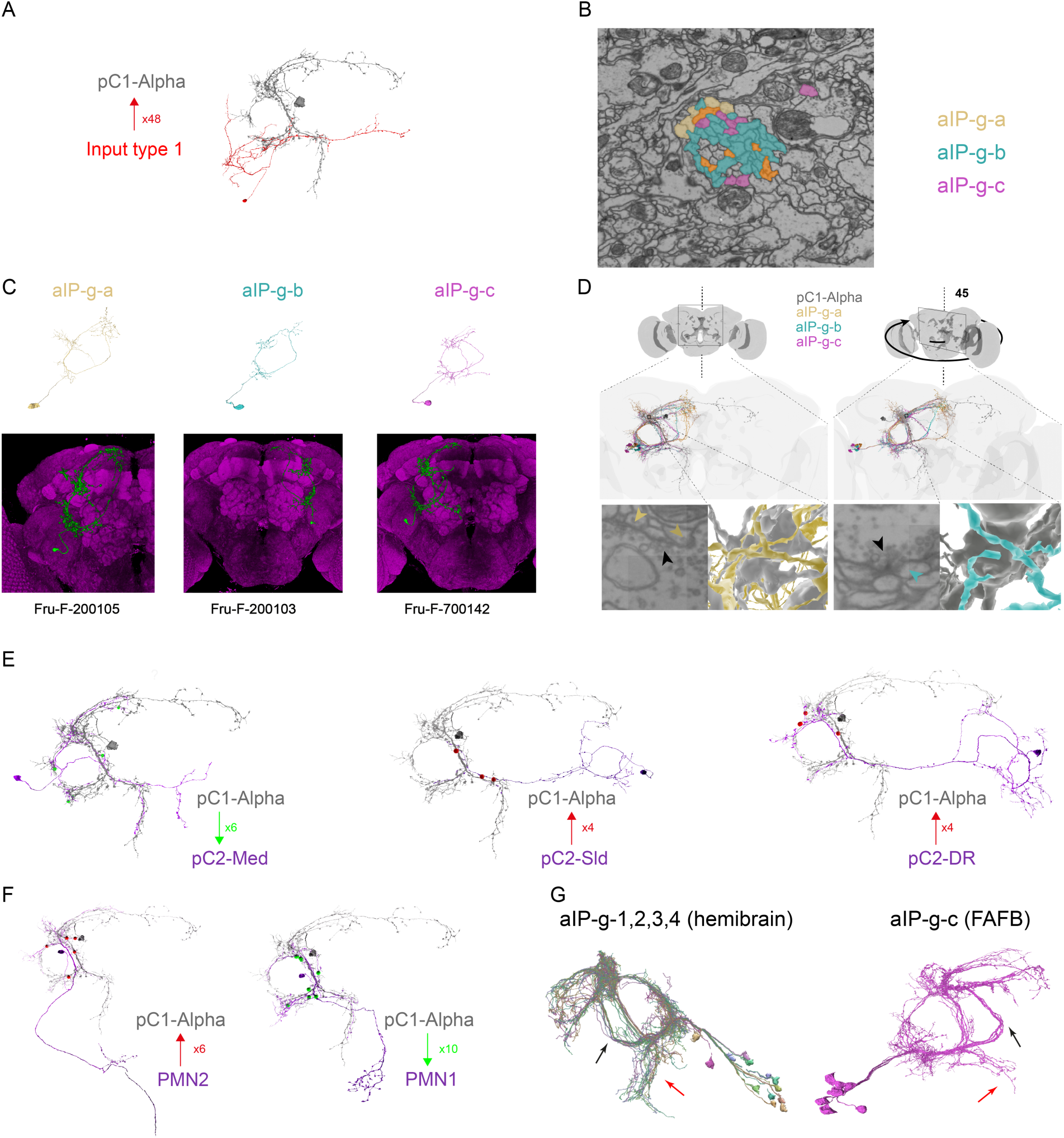
(related to Figure 6) **(A)** The most common neuron of all the neurons presynaptic to pC1-Alpha-l (red). Neurons of this type have a total of 48 presynaptic sites with pC1-Alpha-l (shown in grey). **(B)** A cross section of the bundle that goes out from the aIP-g cell bodies (FlyWire coordinates: 160515, 53776, 3390). aIP-g-a,b,c are colored in yellow, cyan and magenta. One of two cross sections checked in the left hemisphere for aIP-g cell types (second cross section FlyWire coordinates: 161278, 55953, 4590). **(C) Top:** Example aIP-g neurons (one of each type, aIP-g-a,b,c) from FlyWire (following proofreading). **Bottom:** Fru+ clones (from http://www.flycircuit.tw/) that share similar morphology as aIP-g cell types found in FlyWire. **(D)** Example synapses between pC1-Alpha-l and postsynaptic aIP-g-a (left; dyadic synapse) and aIP-g-b (right). **(E)** Example for each of the three pC2-like cell types (purple), each one shown with pC1-Alpha-l (grey) and shared synapses. pC2-Med (Med for medial) cells have a medial projection that overlaps with pC1-Alpha-l medial projection. Some pC2-Med neurons are presynaptic to pC1-Alpha-l and others (as in this example) are postsynaptic. pC2-sld (sld for slide) and pC2-DR are both presynaptic to pC1-Alpha-l. Red dots mark synaptic sites where pC1-Alpha-l is the presynaptic cells. Green dots - pC1-Alpha is postsynaptic. The number of synapses for each example pair are shown. **(F)** Neurons whose morphologies are similar to those of Dsx+ pMN2 neuron (left; presynaptic to pC1-Alpha-l) and Dsx+ pMN1 neuron (right; postsynaptic to pC1-Alpha-l). Red/green dots are synaptic sites, as in (E). **(G)** aIP-g cells in two EM volumetric scans of adult female brains. **Left:** neurons labeled as aIP-g in the hemibrain (Xu et al., 2020) were divided into 4 subgroups (aIP-g-1,2,3,4). **right:** aIP-g-c cells from the FAFB dataset. Black arrows indicate common morphology between aIP-g-c neurons in the two datasets. Black points indicate the point where aIP-g neurons split into aIP-g-a,b,c (see Fig. 6E). Red arrow - a projection that was found in both EM scanned brains, but shows longer projections in the hemibrain. This difference could reflect technical differences or biological variability.

## Supplementary Movies

https://www.dropbox.com/sh/1n8zhumr5bhgbmz/AABAs0lh5MulBKVKJc16P0P8a?dl=0

**Movie S1**

Confidence maps (head in blue, thorax in orange) and part affinity vector fields (white arrows) calculated by LEAP (Pereira et al., 2019) for the male and female. The male has a white painted dot on his back. Male chasing and singing are shown, as well as female shoving. Movie is slowed down 4 times.

**Movie S2**

A sequence of female shoving and chasing. The female is shoving the male while occasionally extending one or two wings, and is then chasing the male while occasionally extending a single wing or contacting the male with her front legs. Finally, the male attempts to copulate, the female spreads her wings and copulation occurs. Movie is in real time. Experimental condition: pC1-Int, d0.

**Movie S3**

Multiple example behaviors: female approaching (Fig. S2C), shoving and circling (Fig. S2D), female headbutting and ‘female mounting’ (Fig. S2E). Following ‘female approaching’ in this example, there is a short epoch of circling. In the ‘shoving and circling’ example, the female is shoving the male before a circling behavior starts (See Fig. S2D, inset). In the ‘female headbutting’ example, the female is extending two wings while headbutting the male, followed by a male jump. In the ‘female mounting’ example, the female is positioning herself behind the male and climbing on his back. Circling and Headbutting examples are from pC1-Int (d0) condition, and female approaching/mounting from pC1-Int (d3) condition.

**Movie S4**

Maximum z-projection (60µm in Z) of the calcium response in a female expressing GCaMP6s pan-neuronaly. Calcium response ((F(t) −Fo/Fo), color coded) is shown 5 minutes before, 5 minutes during and 9.5 minutes after pC1-A activation (using csChrimson). The movie is sped up 20 times.

**Movie S5**

Maximum Z-projection of the calcium response in a female expressing GCaMP6s in Dsx+ cells. pC1 cells in the left hemisphere are shown. Calcium level ((F(t) −Fo/Fo), color coded) is shown 5 minutes before, 5 minutes during and 9.5 minutes after pC1-A activation (using csChrimson). The movie is sped up 20 times.

**Movie S6**

A single pC1-Alpha-l neuron, automatically traced and manually proofread. Inputs (post-synaptic terminals) are shown in red, outputs (pre-synaptic terminals) in green (see also Fig. 6A).

**Movie S7**

pC1 Alpha-l (blue) is shown with example aIP-ga,b,c cells. Synapses are marked in red for inputs (to pC1-Alpha-l) and in green for outputs. Cell type colors (yellow, cyan, magenta) are shown for aIP-g-a,b,c as in Fig. 6F and Supp Fig. S5B-D.

**Movie S8**

pC1-Alpha-l (blue) is shown with neurons that have a similar morphology as known female Doublesex-expressing cells. pC1 subtypes are shown in Fig. 3C and pC2 subtypes are shown in Fig. S5E. pC1-Alpha-l input/output synapses with each example cell are shown in red/green.

